# Pou5f3, SoxB1 and Nanog remodel chromatin on High Nucleosome Affinity Regions at Zygotic Genome Activation

**DOI:** 10.1101/344168

**Authors:** Marina Veil, Lev Yampolsky, Björn Grüning, Daria Onichtchouk

**Affiliations:** Department of Developmental Biology, Institute of Biology I, Faculty of Biology, Albert Ludwigs University of Freiburg, Hauptstrasse 1, 79104, Freiburg, Germany; Department of Biological Sciences, East Tennessee State University, Johnson City, TN, USA; Zoological Institute, Basel University, Basel, Switzerland; Department of Computer Science, Albert Ludwigs University of Freiburg, 79110, Freiburg, Germany; Center for Biological Systems Analysis (ZBSA), University of Freiburg, Freiburg, Germany; Signalling Research centers BIOSS and CIBSS, Freiburg, Germany; Institute of Developmental Biology RAS, Moscow, Russia

## Abstract

The zebrafish embryo is mostly transcriptionally quiescent during the first 10 cell cycles, until the main wave of Zygotic Genome Activation (ZGA) occurs, accompanied by fast chromatin remodeling. At ZGA, homologs of mammalian stem cell transcription factors (TFs) Pou5f3, Nanog and Sox19b bind to thousands of developmental enhancers to initiate transcription. So far, how these TFs influence chromatin dynamics at ZGA has remained unresolved. To address this question, we analyzed nucleosome positions in wild-type and Maternal-Zygotic (MZ) mutants for *pou5f3* and *nanog* by MNase-seq. We show that Nanog, Sox19b and Pou5f3 bind to the High Nucleosome Affinity Regions (HNARs). HNARs are spanning over 600 bp, featuring high *in vivo* and predicted *in vitro* nucleosome occupancy and high predicted propeller twist DNA shape value. We suggest a two-step nucleosome destabilization-depletion model, where the same intrinsic DNA properties of HNAR promote both high nucleosome occupancy and differential binding of TFs. In the first step, already prior to ZGA, Pou5f3 and Nanog destabilize nucleosomes on HNAR centers genome-wide. In the second step, post-ZGA, Nanog, Pou5f3 and SoxB1 maintain open chromatin state on the subset of HNARs, acting synergistically. Nanog binds to the HNAR center, while the Pou5f3 stabilizes the flanks. The HNAR model will provide a useful tool for genome regulatory studies in the variety of biological systems.

## Introduction

The development of multicellular organisms is first driven by maternal products and occurs in the absence of transcription (Tadros and Lipshitz 2009). The main wave of the embryo’s own transcription, Zygotic Genome Activation (ZGA), occurs several hours after the egg is fertilized. The mechanistic reasons for this transcriptional delay are not completely understood. In *Drosophila* and zebrafish *Danio rerio*, transcription after ZGA depends on a small number of maternal enhancer-binding Transcription Factors (TFs). Zygotic genome activators include Zelda in *Drosophila* (Liang et al. 2008), and three homologs of pluripotency TFs, Pou5f3, Nanog and Sox19b in zebrafish (Lee et al. 2013; Leichsenring et al. 2013). Recent analysis of chromatin accessibility and gene expression also implicated transcription factors POU5F1 and Nfya as ZGA activator in humans (Gao et al. 2018) and mice (Lu et al. 2016), respectively. The interactions of ZGA activators with chromatin are only starting to be understood, and it is unclear if the same ZGA mechanisms are implemented in different animals (Winata et al. 2018).

The basic units of chromatin structure are the nucleosomes, which restrict the *in vivo* access of most TFs to their target sites (Beato and Eisfeld 1997; Luo et al. 2014). While many TFs cannot bind their target site in the context of nucleosome DNA *in vitro*, cooperative interactions among multiple factors allow binding, even in the absence of defined orientation of their recognition motifs in DNA (Adams and Workman 1993, Zinzen et al. 2009). In steady-state cell culture systems, most of the TF binding sites fall within nucleosome-free DNA regions (Thurman et al. 2012), and it is difficult to conclude if the chromatin accessibility is a cause or a consequence of TF binding. Studies of hematopoietic cell fate transitions suggested a hierarchical model, where a relatively small set of “pioneer” TFs collaboratively compete with nucleosomes to bind DNA in a cell-type-specific manner. The binding of pioneer TFs, also called lineage-determining factors and master regulators, is hypothesized to prime DNA by moving nucleosomes and inducing the deposition of epigenetic enhancer marks. This enables concurrent or subsequent binding of signal-dependent transcription factors that direct regulated gene expression (Heinz et al. 2010; Trompouki et al. 2011; Li et al. 2018). Two alternative scenarios of TF-guided chromatin opening at enhancers during cell-fate transitions have been suggested: the nucleosome-mediated cooperativity model, and the model based on specialized properties of pioneer transcription factors (Calo and Wysocka 2013; Slattery et al. 2014). The nucleosome-mediated cooperativity model postulates that if the recognition motifs for multiple cooperating TFs are located within the length of one nucleosome, the nucleosome will be removed (Mirny 2010; Moyle-Heyrman et al. 2011). The alternative model evokes special properties of pioneer factors to bind their DNA sites directly on nucleosomes (Iwafuchi-Doi and Zaret 2016). The pioneer factor model predicts that pioneer TFs will engage into the silent chromatin, while the majority of non-pioneer TFs will occupy nucleosome-depleted regions.

In *Drosophila*, the ZGA activator Zelda binds to regions of high predicted nucleosome occupancy and creates competency for other factors to bind the DNA, thus providing an example of a pioneer factor (Schulz et al. 2015; Sun et al. 2015). Genomic binding of zebrafish ZGA activator Pou5f3 is detectable at 512-cell stage, just prior to the main ZGA wave (Leichsenring et al. 2013). Binding of Pou5f3 to the enhancers of its early regulated target genes could be significantly outcompeted by increasing histone concentration at ZGA (3 hpf, 10^th^ cell cycle, Joseph et al. 2017). In zebrafish, nucleosome positioning signals guide the transcription start site (TSS) selection after ZGA, but not before, suggesting that the rules of nucleosome positioning may change over ZGA (Haberle et al., 2014).

Genetic ablation of maternal and zygotic expression of Pou5f3 and Nanog results in severe pleiotropic phenotypes, global deregulation of transcription, and gastrulation arrest (Lunde et al. 2004; Onichtchouk et al 2010; Gagnon et al. 2018; Veil et al. 2018), suggesting that these factors act at the genome-wide level. However, it is not clear whether Pou5f3 or Nanog preferentially target the regions occupied by nucleosomes prior to ZGA or are able to remove nucleosomes from the regions they bind to. We address these questions below.

## Results

### The properties of Transcription Factor Binding Sites (TFBS) bound by Pou5f3, SoxB1 and Nanog individually or in combination

Our analysis strategy was to categorize the genomic regions by their binding of Pou5f3 (P), SoxB1 (S), and Nanog (N), individually or in combination. For this analysis, we focused on the transcription factor binding sites (TFBS) within ChIP-seq peaks from our own and others’ previous work (Leichsenring et al. 2013, Xu et al. 2012), and on the group of randomly chosen control regions of similar size. Based on TFs binding data, genomic sites were classified into the following groups: P and N groups (Pou5f3 and Nanog bind individually), PS, PN, and SN groups (TFs bind pair-wise), and the PSN group (all three TFs bind). The group of genomic regions occupied by Pou5f3 alone and only pre-ZGA, termed Ppre, served as an additional control. We set to compare the relative changes in nucleosome occupancy using MNase-seq method for each of these groups between MZ*spg*, MZ*nanog* mutants and the wild-type. In the cases where we found statistically significant differences in nucleosome occupancy between the groups, we evaluated to which extent these differences could be explained by sequence-specific binding of the TFs to their consensus recognition motifs. We use the terms “consensus binding” and “nonconsensus binding”, as suggested by Afek et al. (2015), to distinguish between TF binding to its cognate consensus motifs on DNA (“consensus binding”), and TF binding at the absence of motifs (“nonconsensus binding”). The heat maps of TF occupancy for the seven groups are shown in Fig. 1A, summary profiles in Fig. S1, the genomic regions are listed in the Table S1. We performed *de-novo* motif search to estimate the relative abundance of the sequence-specific recognition consensus motifs for each TF in each group. In addition to the sequences closely matching known Pou5f3, SoxB1 and Nanog consensus motifs (*pou:sox*, *sox*, *nanog1* and *nanog2*), we detected enrichment for generic motifs for C2H2-type Zinc-finger proteins and beta-HLH domain transcription factors, dinucleotide repeats, and tetranucleotide ATSS repeats on the regions bound by Pou5f3 alone. Fig. 1B shows distribution of the motifs within the TFBS groups (see Fig. S2 for all logos and Table S2 for matrices).

**Figure 1.**
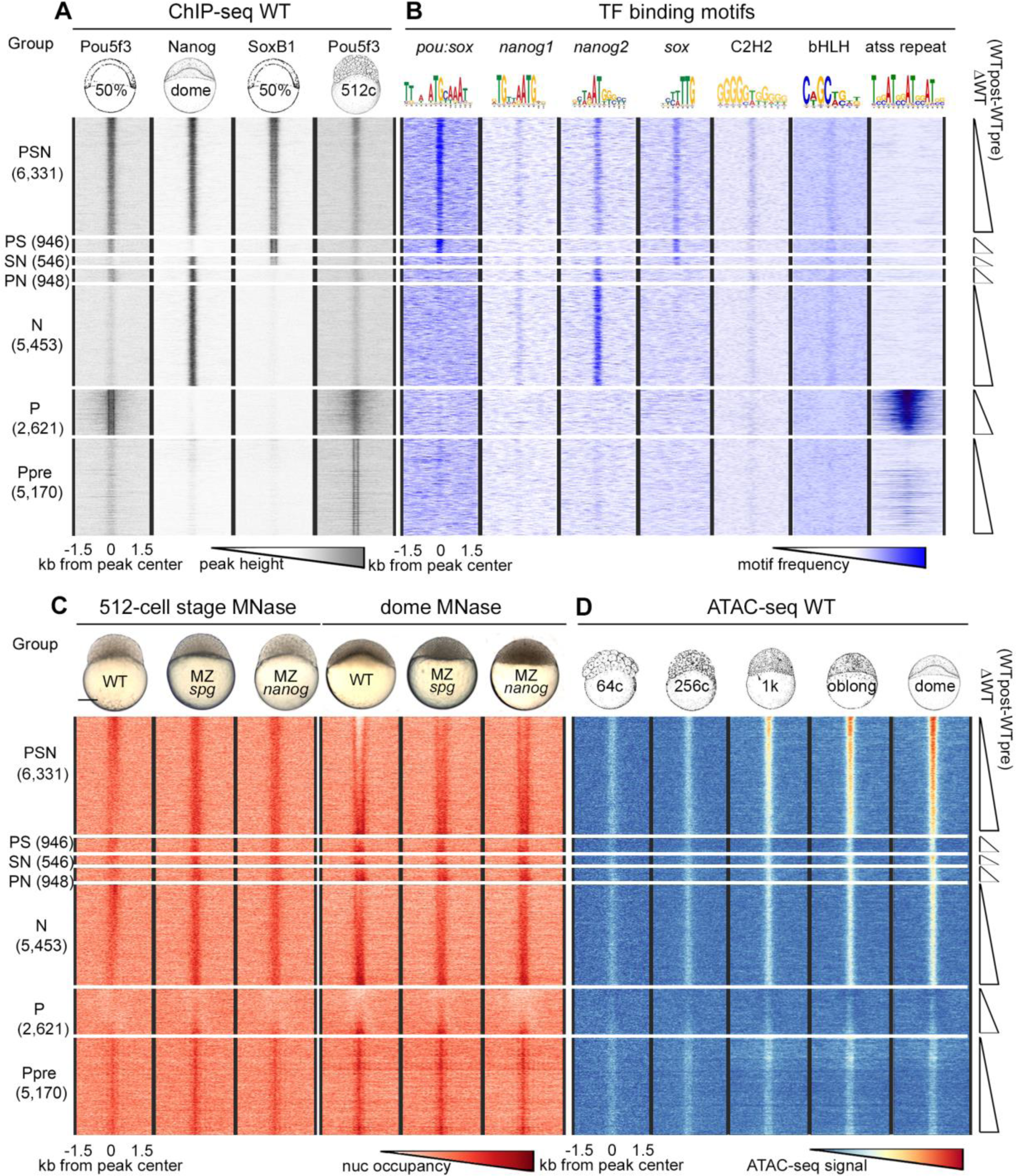
Nucleosomes cover all TFBS pre-ZGA, and are depleted from PSN triple-bound regions post-ZGA. Heat maps of 6 groups of genomic regions defined by combinatorial binding of Pou5f3, Nanog and SoxB1 post-ZGA, as indicated at the left, and Ppre group bound by Pou5f3 only pre-ZGA. 3 kb genomic regions were aligned at the ChIP-seq peak centers. Within each group, the data were sorted by ascending difference between wild-type (WT) post-ZGA and pre-ZGA nucleosome occupancy (∆WT post-pre). (*A*) TF binding (Pou5f3 50% epiboly, Nanog and SoxB1 dome stage, Pou5f3 512-cell stage) (*B*) Occurrence of TF-binding motifs. (*C*) Nucleosome occupancy in the embryos of indicated genotypes. (*D*) Accessible chromatin signals in pre-ZGA (64c, 256c, c – cell stage), ZGA (1K) and post-ZGA stages (oblong, dome). Scale bar (C) indicates 200 µm.

Pou5f3, SoxB1 and Nanog preferentially bind to the enhancers of developmental genes (Xu et al. 2012; Leichsenring et al. 2013). To compare the enrichment in developmental enhancers in the groups defined above, we used GREAT analysis (McLean et al. 2010). The enrichment in enhancers within PSN group was an order of magnitude higher than that in N, PS, PN, SN, Ppre groups (Fig. S3). We found no enrichment for P group, indicating that nonconsensus binding of Pou5f3 alone post-ZGA does not mark enhancers.

### All TFBS are occupied by nucleosomes before ZGA; nucleosomes are depleted only from the triple-bound Pou5f3, Nanog and SoxB1 regions after ZGA

We isolated chromatin from the wild-type, MZ*spg* and MZ*nanog* mutants at pre-ZGA (512-cell stage) and post-ZGA (dome stage) and performed micrococcal nuclease treatment followed by deep sequencing of the resulting fragments (MNase-seq) as described (Zhang et al. 2014, Fig. S4, Table S3). At the 512-cell stage, all TFBS had higher nucleosome occupancy than surrounding sequences (Fig. 1C, 512-cell stage). At dome stage, a nucleosome-depleted region appeared in the PSN group, with these depletions being strongly reduced in the MZ*nanog* and MZ*spg* mutants compared to the wild-type (Fig. 1C, dome). Single or dual Pou5f3, SoxB1 or Nanog occupation sites remained covered by nucleosomes. We compared our data with recently published ATAC-seq based chromatin accessibility data (Liu et al. 2018). Weak ATAC-seq signals which did not meet the standard cutoffs in Liu et al. 2018 and Meier et al. 2018 were detectable when aligned on all TFBS (Fig. 1D). Before ZGA, the elevated nucleosome occupancy (MNase-seq) and accessible chromatin (ATAC-seq) signals colocalized on all TFBS irrespectively to their regulatory potential (Fig. 1C, 512-cell stage, Fig. 1D, 64-and 256-cell stage). We interpreted mixed signals of opposite direction as local destabilization of nucleosomes on TFBS. Mixed high MNase and high ATAC-seq signals could reflect heterogeneity of the cells in the embryo: in a few cells TFBS was nucleosome-free, while in most of the cells TFBS was covered by nucleosome. Low MNase and high ATAC-seq signals colocalized only on PSN group, indicating nucleosome depletion from only PSN regions post-ZGA. All other TFBS were covered by destabilized nucleosomes, pre-and post-ZGA.

### Pou5f3 and Nanog non-specifically reduce nucleosome occupancy before ZGA and act sequence-specifically after ZGA

We tested whether the differences in pre-and post-ZGA nucleosome occupancy between each of the two mutants, MZ*spg* and MZ*nanog* and the wild-type were significantly different from 0 in each TFBS group and significantly different among the TFBS groups. This was the case for both stages (Fig. S5, Table S4 for 1-way ANOVA for all comparisons). At the 512 cell stage, MNase-seq signal in both mutants was higher than in the wild-type for all TFBS (except for P group in MZ*nanog*, Fig. S5B,C). At dome stage, MNase-seq signal in both mutants mostly increased in the PSN group compared to the wild-type (Fig. S5,D-F). This indicated that Pou5f3 and Nanog displace nucleosomes at both stages. To test if nucleosome displacement by TFs correlates with gene expression, we linked each TFBS to the nearest promoter defined in Haberle et al. (2014), and divided them into four categories (early zygotic, late zygotic, maternal-zygotic and non-expressed, Fig. S6A). Nucleosome displacement by TFs pre-ZGA, and in double-and single-TF-bound TFBS post-ZGA did not differ between gene expression categories. On the PSN (triple-bound) TFBS linked to the early zygotic genes, nucleosome displacement post-ZGA was stronger and more dependent on Pou5f3 and Nanog, than on the PSN TFBS linked to the other categories (Fig. S6B-D).

To test whether the same regions within the PSN group are affected at the 512-cell and dome stages, we ranked 6,331 PSN regions by descending difference between pre-and post-ZGA nucleosome occupancy (ΔWT=WTpost-WTpre, Fig. 1C,D) and divided them to octiles O1-O8, O1 being the most open at dome stage. The order of octiles by nucleosome occupancy did not match between the 512-cell and dome stages (Fig. 2A,B); neither did the mutant effects (Fig. S7A,B). The ranking of octiles at dome stage correlated with the ATAC-signal strength starting from ZGA (1K, Fig. 2C) and with the occupancy by Pou5f3 and Nanog TFs (Fig. 2D). The reduction in nucleosome occupancy by Pou5f3 and Nanog assayed by the MNase method at post-ZGA dome stage corresponded to the increase of the accessible chromatin signals assayed by ATAC-seq at post-ZGA oblong stage (Pou5f3 and Nanog morpholino knockdown experiments of Liu et al. 2018, Fig. S7B,C).

**Figure 2.**
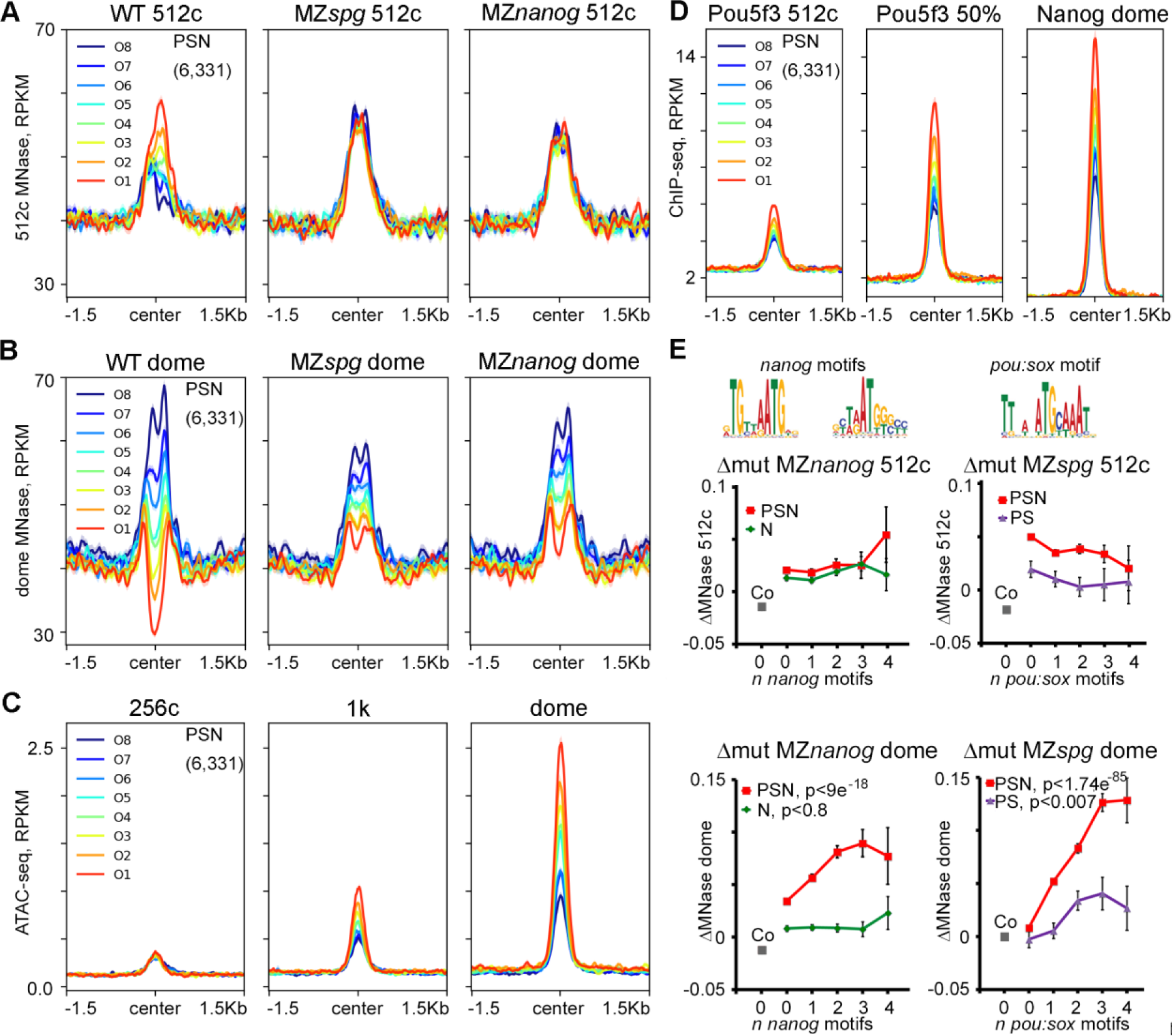
Pou5f3 and Nanog displace nucleosomes from different regions before and after ZGA. (*A-D*) PSN group regions ranked into octiles by ascending difference between WT post-ZGA and pre-ZGA MNase signal (∆WT post-pre). 3 kb genomic regions were aligned at the ChIP-seq peak centers. Summary plots per octile (*A*) nucleosome occupancy 512-cell stage, (*B*) nucleosome occupancy dome, (*C*) ATAC-seq, (*D*) TF occupancy. (*E*) TF effects on nucleosome density were estimated as MNase signal difference between the wild-type and indicated mutants (∆mut) and related to the number of specific motifs (0, 1, 2, 3 and ≥ 4 motifs) per 320 bp binding region. Motifs are indicated on top. Pou5f3 and Nanog act non-sequence specifically at 512 cell stage (upper row). At dome stage (lower row), Pou5f3 and Nanog act by sequence-specific binding to their motifs on triple-occupancy PSN regions, (PSN – red), but not on single and double TFBS (N - green, PS - violet). Co – random control regions (0 motifs), grey. Y-axis – ∆mut.

To evaluate the contribution of TF consensus binding to the nucleosome displacement, we compared MNase signal changes between each mutant and the wild-type (∆mut MZ*spg* and ∆mut MZ*nanog*) within PSN, N or PS groups on the regions with 0, 1, 2, 3 and ≥ 4 cognate Pou5f3/SoxB1 or Nanog motifs (Fig. 2E, Table S5 for 1-way ANOVA). ∆mut MZ*spg* and ∆mut MZ*nanog* did not depend on the number of motifs at the 512-cell stage (Fig. 2E, 512c). However, ∆mut MZ*spg* and ∆mut MZ*nanog* on TFBS without motifs was higher than in randomly chosen control sequences, suggesting either indirect or non-specific effects of the TFs (Fig. 2E, 512-cell stage, grey Co data point, see Table S6 for Student’s *t*-test). At the dome stage, ∆mut MZ*spg* and ∆mut MZ*nanog* depended on the number of respective motifs in PSN group regions, but not in N group and only marginally in PS group (Fig. 2E, dome). These results paralleled accessible chromatin changes assayed by morpholino knockdown experiments and ATAC-seq (Fig. S7D). Thus, in contrast to non-specific pre-ZGA events, post-ZGA nucleosome depletion relied on specific binding of Pou5f3, Nanog and SoxB1 (PSN) and required synergistic action of all three TFs.

### Pou5f3 and Nanog bind chromatin independently from pre-existing chromatin modifications and DNA methylation

The mixed elevated MNase and ATAC-seq signals observed on all TFBS could reflect nucleosome destabilization by either distinct sequence composition or epigenetic pre-marking. Murphy et al. (2018) have described “placeholder” H2AZ/H3K4me1 containing nucleosomes, which occupy the regions lacking DNA methylation in cleavage embryos. Only 5.3% of TFBS defined in our study overlapped with placeholder nucleosome locations (Fig. S8A), and these 5.3% TFBS were bound by Pou5f3 and Nanog similarly to the rest of the sites (Fig. S8B-D). Liu et al. (2018) suggested that Pou5f3 and Nanog specifically recognize methylated motifs; we did not find a support for this statement (see Fig. S9 and legend). As we could not find strong evidence for pre-existing epigenetic marking, we focused on sequence composition of TFBS regions, in order to explain nucleosome destabilization effects.

### Pou5f3, SoxB1 and Nanog bind to 600 bp long asymmetric High Nucleosome Affinity Regions (HNARs)

The reasons for high nucleosome occupancy of pre-ZGA TFBS could be intrinsic nucleosome-favoring DNA features. Nucleosomes cover 75-90% of the genome (Kornberg 1974) and can form virtually anywhere, but the properties of the underlying DNA may change nucleosome affinity up to several orders of magnitude (Field et al. 2008). Increased GC content and 10 bp periodic fluctuations of dinucleotides underlie the high ability of DNA to wrap around histone octamers, while AT-rich sequences strongly disfavor nucleosome formation (Satchwell et al. 1986; Field et al. 2008; Chung and Vingron 2009; Kaplan et al. 2009; Tillo and Hughes 2009). To evaluate if Pou5f3, SoxB1, and Nanog preferentially recognize nucleosome-positioning signals, we used nucleosome predictions based on *in vitro* nucleosome DNA sequence preferences mentioned above (Kaplan et al. 2009). The predicted nucleosome occupancy is scored by a probability for each nucleotide position to be located in the nucleosome center (or dyad). We selected the maximal [nucmax] score (for the most likely predicted dyad position) within the mean length of ChIP-seq peak (320 bp) for each TFBS and control region and compared them across the groups. [Nucmax] predicted scores of all (individual, double-and triple-bound) TFBS were higher than respective scores for random control regions (Fig. S10A). Notably, the [nucmax] scores of TFBS groups where Nanog binds (i.e., N, PN, SN, and PSN groups) were significantly higher than of those of the groups where Nanog does not bind (Fig. S10B, Table S7).

The minimum nucleosome prediction scores within the 320 bp interval [nucmin] most likely corresponded to the predicted inter-nucleosomal sequences within the region (Segal et al. 2006). In all TFBS [nucmin] scores were also significantly higher than the respective scores for random control regions (Fig. S10C,D, Table S8). The difference in [nucmin] predictions agreed with experimental data: experimental nucleosome occupancy of random control regions aligned on [nucmin] positions was below the genomic average, as predicted, but it was above the genomic average in [nucmin] positions of all TFBS (Fig. S10E-L). These data implied that high nucleosome affinity regions underlying TFBS span at least 320 bp, i.e. more than two nucleosome lengths.

TF recognition motif sequences may themselves contain high predicted nucleosome occupancy features, or, alternatively, the high predicted nucleosome occupancy may be encoded by sequences neighboring the recognition motifs. To distinguish between these possibilities, we compared predicted nucleosome occupancy of the motifs that are actually bound by Nanog or Pou5f3/SoxB1 (within 320 bp of respective TFBS), with predicted nucleosome occupancy of randomly occurring unbound motifs (within the interval from 1 to 1.5 kb away from the TFBS center). All bound motifs could be discriminated from the unbound ones by higher predicted nucleosome occupancy of neighboring sequences (Fig. 3A-C). Nucleosome prediction profiles around *nanog1* and *nanog2* motifs were higher and different in shape from the profiles on *pou:sox* motifs (compare Fig. 3A,B with Fig. 3C), while *pou:sox* motif profiles (Fig. 3C) were similar to the flattened profiles on the regions bound by Pou5f3 pre-ZGA (Ppre, Fig. 3D).

**Figure 3.**
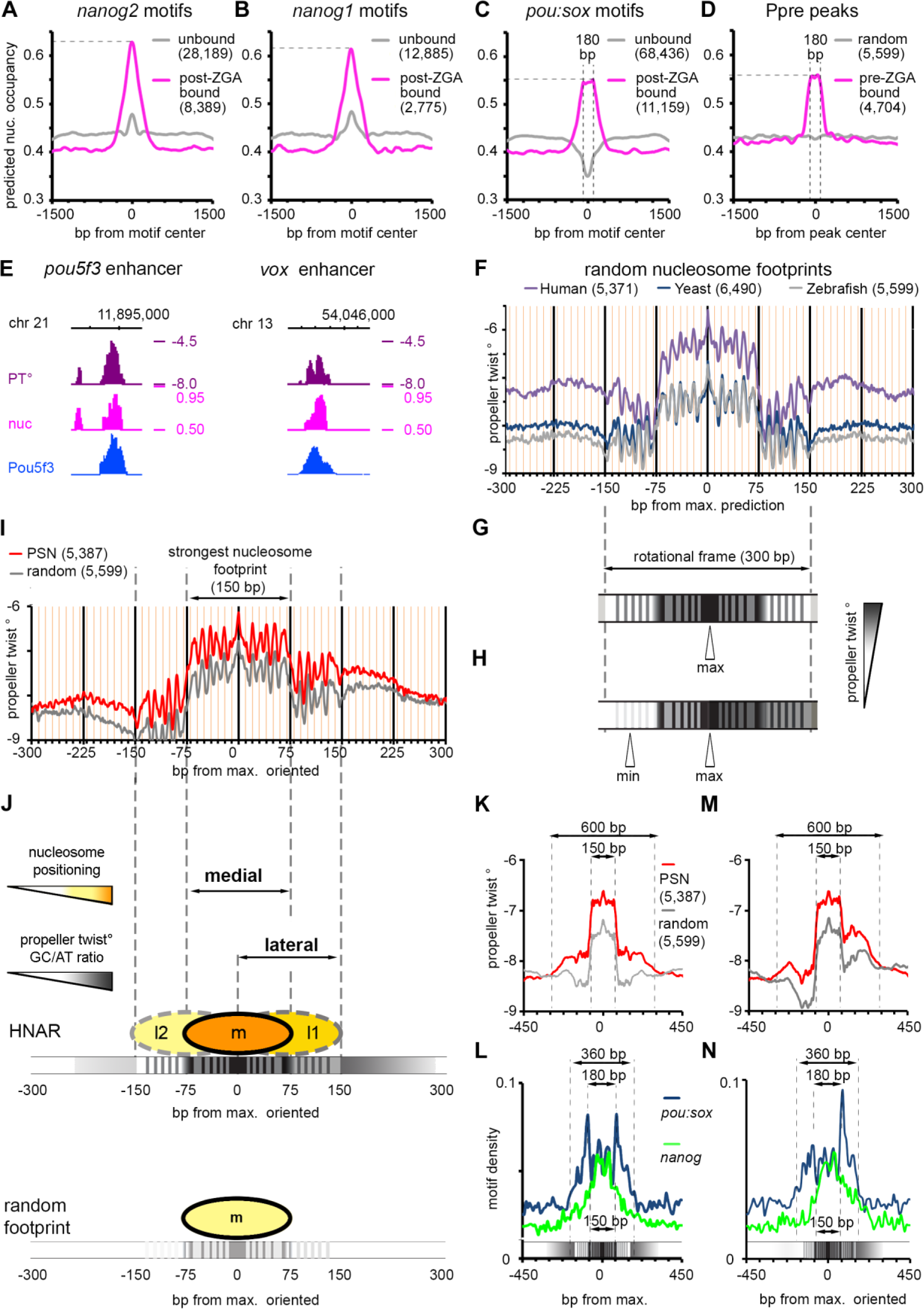
Pou5f3, SoxB1 and Nanog bind to the High Nucleosome Affinity Regions (HNARs). (*A-C*) Mean predicted nucleosome occupancy plots on the indicated motifs bound by respective TF (magenta) versus unbound random motif matches (gray). (*D*) Mean predicted nucleosome occupancy plot centered on Pou5f3 pre-MBT TF binding peaks (magenta) versus random sequences (gray). (*E*) Pou5f3 binds to the regions of high predicted nucleosome occupancy and Propeller Twist on the enhancers of *pou5f3* and *vox*. PT-propeller twist (°), nuc -*in vitro* nucleosome occupancy prediction, tick marks every 1 kb. (*F*) Propeller twist (PT) summary plot of random zebrafish, yeast, and human genomic regions aligned at the base pair with maximal predicted nucleosome occupancy within 320 bp [nucmax]. Yellow X-axis gridlines 10 bp apart. (*G*) Scheme of the PT periodic oscillations in 300 bp around [nucmax]. (*H*) Scheme of the oriented PT plot aligned at [nucmax]. (*I*). Oriented PT plot of PSN group (red) and control genomic regions (gray). (*J*). Scheme of medial and lateral strong nucleosome footprints on HNAR. Central 300 bp periodic frame supports alternative medial, lateral 1 and lateral 2 nucleosome positions with decreasing strength (m>l1>l2=random footprint); in random sequence only medial footprint is supported (bottom). (*K,M*) Symmetric (K) and oriented (M) smoothened PT plots (80 bp moving average) of PSN and control genomic regions, aligned at [nucmax]. (*L,N*) Distribution of *nanog* and *pou:sox* motifs in the PSN group for (K) and (M), respectively. Motif density in bp motif per bp sequence.

There is no agreement in the literature on the intrinsic nucleosome occupancy features within regulatory sequences. Under-representation of nucleosome positioning sequences in enhancers has been reported (Daenen et al. 2008; Papatsenko et al. 2009; Khoueiry et al. 2010; He et al. 2010), while other studies reported their over-representation (Tillo et al. 2010; Barozzi et al. 2014; Sun et al. 2015). Our data strongly support the notion that intrinsic nucleosome occupancy in zebrafish early enhancers is high, but in the view of the contradictory literature we aimed to confirm this conclusion by using an independent approach. Propeller twist (PT), the angle between the heterocycles in the two complementary bases in a base pair, is a DNA shape parameter that is not directly encoded by primary sequence. PT values have been related to the flexibility of DNA bending around proteins (el Hassan and Calladine 1996) and strongly correlate with the ability of DNA to wrap around nucleosomes in yeast (Lee et al. 2007, Gan et al. 2012). Intrinsic nucleosome-forming preferences of the DNA on the known PSN-bound enhancers of *pou5f3* (Parvin et al. 2008), *vox,* and *vent* genes (Belting et al. 2011) were independently captured by the primary sequence–derived model of Kaplan et al. (2009) and high values of propeller twist (Fig. 3E, Fig. S11).

We aimed to further describe the length and structure of putative enhancers which underlie TFBSs by combining propeller twist (PT) and the nucleosome predictions based on Kaplan et al. (2009) model. Since it is unknown how the features captured by sequence-based nucleosome positioning programs are reflected in PT values, we started from the control sequences. We randomly selected 5,000-6,000 genomic sequences from human and yeast genomes as 320 bp control sequences and found [nucmax] predicted nucleosome scores (dyad) within each region by the model of Kaplan et al. (2009). Then we plotted the average PT values for zebrafish, human and yeast aligned on the [nucmax] position. As shown in Fig. 3F, a sharp increase in PT marked the central 150 bp nucleosome footprint, which was embedded into a 300 bp periodic shape. The differences in the height of PT graphs roughly corresponded to the median GC content in zebrafish, yeast, and human genomes (36.8%, 38.4% and 40.9%, respectively) as provided by NCBI.

The ~10 bp periodicity for PT prediction within the central 300 bp of DNA (Fig. 3G) represents a single rotational frame, which was reported to stabilize nucleosome positioning on DNA (see Fig. S12 and legend). To uncover possible hidden asymmetry of the nucleosome positioning signals we oriented each 320 bp genomic region centered on [nucmax] so that the minimum nucleosome prediction value within +/-160 bp around [nucmax] is positioned at the left (5’) side of the interval, as shown on Fig. 3H. We compared summary PT heatmaps of TFBS groups aligned on [nucmax] or [nucmin] with their nucleotide composition and found that higher PT values corresponded to increased GC content (Fig. S13A,B, the legend). Thus, PT sensitively reflected two attributes of DNA affinity to nucleosomes: GC content and dinucleotide rotational periodicity. We next used the [nucmax] and [nucmin] positions as viewpoints to estimate the differences between TFBS and random nucleosome footprints using symmetric and asymmetric PT plots. All TFBSs except P and PS groups differed from the control group by higher propeller twist values over ~600 bp around [nucmax], as shown in Fig. 3I for PSN group regions (see Fig. S14-15 for all groups). We define here the 600 bp DNA region with high predicted nucleosome occupancy/propeller twist/GC content, which is centered on a 300 bp periodic frame, as a High Nucleosome Affinity Region (HNAR).

### HNAR structure supports two strong nucleosome positioning sequences, medial and lateral

The random nucleosome footprint and HNAR, both oriented from low to high nucleosome prediction/PT, are shown in Fig. 3J. PT values/relative GC content is higher in HNAR, which should theoretically result in stronger nucleosome positioning. The graph above HNAR shows three positions within the 300 bp frame, given that high PT and periodic PT signal represent two independent factors that stabilize nucleosome on DNA. The central (medial), lateral 1 and lateral 2 positions are supported by 10 periodic peaks each. The medial position (m) is the most stable, the l1 (lateral) position is weaker, because of the overall lower PT, but is still stronger than random nucleosome footprint. The l2 position is the weakest, being supported by even lower PT value compared to m and l1 positions. In the random nucleosome footprint only the medial position is stabilized by underlying DNA features (Fig. 3J bottom). TF-binding motifs for Nanog and Pou5f3/SoxB1 were differentially distributed around HNAR center. When HNAR was oriented symmetrically (as shown in Fig. 3K), most *nanog* motifs localized within the medial 150 bp nucleosome footprint, while *pou:sox* motifs were present over the whole 300 bp central periodic shape, with two peaks 180 bp apart from each other (Fig. 3L). In the oriented projection of HNAR (Fig. 3M) the *pou:sox* motifs added up in one peak within the higher PT flank (lateral footprint), while the *nanog* motifs were mostly central (Fig. 3N). The nucleosome distribution in the model on Fig. 3J is theoretical, and we next addressed if it could help us to interpret *in vivo* data.

### Pou5f3 and Nanog non-specifically destabilize nucleosomes on HNAR centers pre-ZGA

Before ZGA, Pou5f3 and Nanog destabilized nucleosomes on all TFBS groups, independently from the presence of their consensus motifs. Could structural properties of HNARs explain this effect? To find out if it is the case, we divided the TFBS and random control regions, matched by [nucmax] value, into quartiles Q1-Q4, Q1 having the lowest [nucmax] value. The PT and nucleosome prediction graphs for control oriented nucleosome footprints are shown in Fig. 4A,B (see Fig. S16 A,B for the TFBS groups). High MNase and high ATAC-seq signals co-localized within random strong nucleosome footprint (Fig. 4C,D), indicating that footprint features alone caused destabilization of nucleosomes. On random nucleosome footprint and on TFBS, ATAC-seq signal peaked on the predicted dyad (Fig. 4D), while nucleosomes were shifted to the lateral positions (black arrows on Fig. 4C). The strength of the ATAC-seq signal positively correlated with the predicted nucleosome occupancy value in all stages and groups (Fig. 4D, Fig. S16 C,D for TFBS groups and Table S9 for statistics). Thus, HNAR structure itself promoted nucleosome destabilization, employing a mechanism which relies on predicted nucleosome occupancy. We tested if the nucleosome displacement effects of Pou5f3 and Nanog pre-ZGA contribute to this mechanism. Pre-ZGA, Pou5f3 and Nanog displaced nucleosomes from the random nucleosome footprints and HNAR centers, and the strength of displacement increased with the predicted nucleosome occupancy (Fig. 4 E,F, Fig. S17A-D). Pou5f3 ChIP-seq signals at 512-cell stage were present on all HNAR centers and even on control footprint (Fig. S17E), indicating that Pou5f3 may act by direct binding. At 512-cell stage, the nucleosome displacement by Pou5f3 and Nanog increased with the predicted nucleosome footprint strength in all TFBS groups and the control (Fig. 4G shows the whole data set, Fig. S18 shows TFBS groups, Table S10 shows statistics). Post-ZGA, nucleosome displacement only by Nanog depended on nucleosome prediction strength (Fig. 4G), suggesting that Pou5f3 acts differently before and after ZGA. We concluded, that non-specific nucleosome destabilization by Pou5f3 and Nanog pre-ZGA depends on HNAR structural features, is not restricted to enhancers and occurs genome-wide.

**Figure 4.**
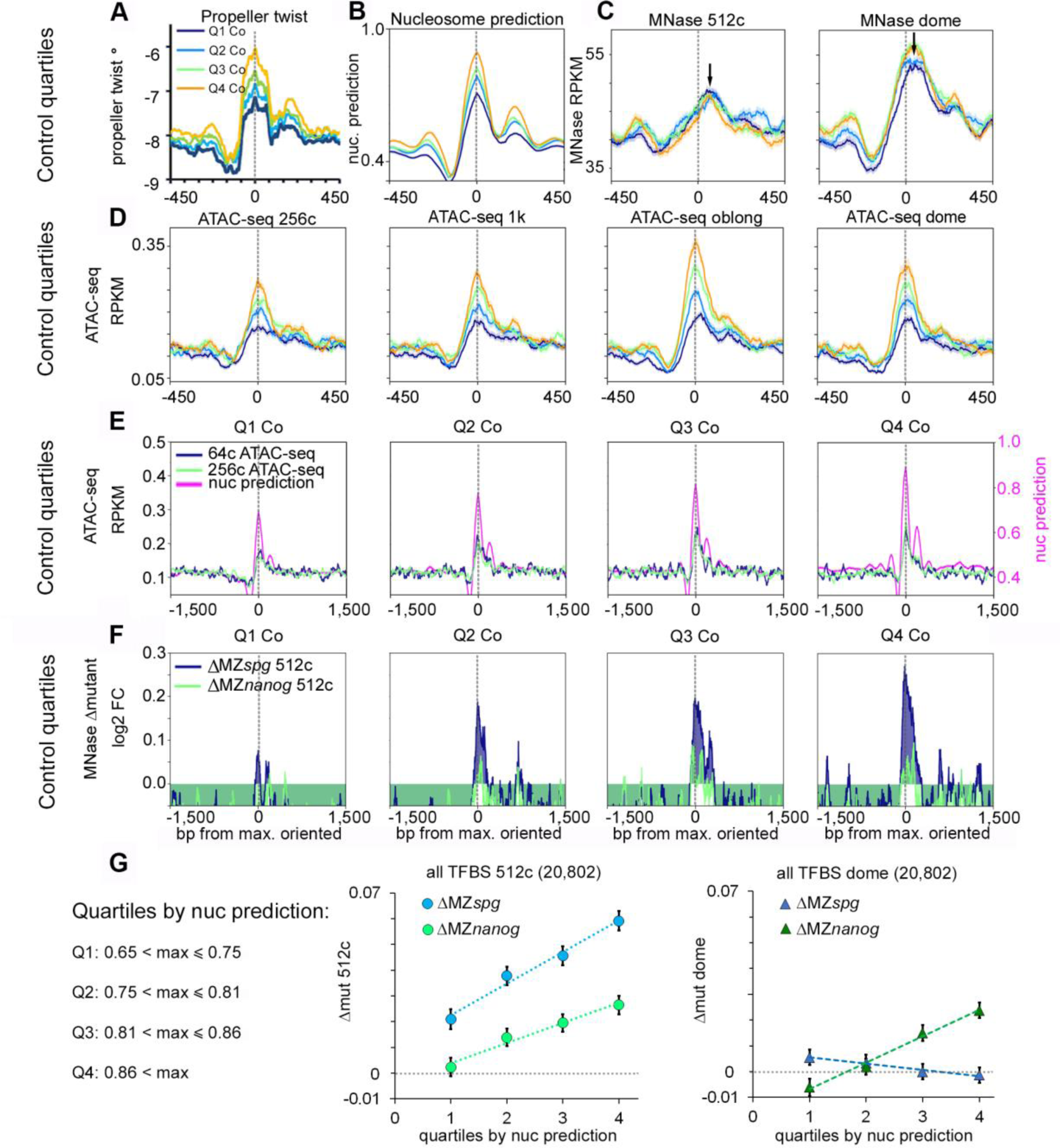
Pou5f3 and Nanog randomly displace nucleosomes from HNAR centers pre-ZGA. (*A-D*) Random control genomic regions were ranked by ascending nucleosome prediction score into quartiles Q1-Q4, aligned on HNAR centers ([nucmax], gray dotted lines), and oriented as in Fig. 3H. *(A)* PT° *(B)* nucleosome prediction *(C)* MNase, black arrows show the lateral shift from [nucmax] *(D,E)* ATAC-seq signal colocalizes with [nucmax] and increases with nucleosome prediction value. (*F*) Random control quartiles: nucleosome displacement from HNAR centers by Pou5f3 and Nanog at 512 cell stage increases with nucleosome prediction value. Gray dotted lines - [nucmax]. (*G*) Dependencies of Δmut (nucleosome occupancy) from nucleosome prediction strength in the whole TFBS data set. 512-cell stage – left, dome – right.

### Sequence-specific Nanog binding at the HNAR center and Pou5f3 binding on +90 bp lateral position deplete nucleosomes post-ZGA

Widespread nucleosome destabilization may facilitate specific binding of TFs, if their motifs are present on HNAR centers. Nucleosome-mediated cooperativity model (also referred as “assisted loading” or “collaborative binding” (Spitz and Furlong, 2012)) postulates that nucleosome depletion occurs when multiple TFs specifically bind their motifs on DNA within one strong nucleosome positioning sequence. HNAR contains at least two overlapping strong nucleosome positioning sequences, medial and lateral footprints (Fig. 3J). Our next question was if synergistic nucleosome depletion by Pou5f3, SoxB1, and Nanog after ZGA depends on the amount and distribution of the specific TF motifs on medial (−75 to 75 bp from HNAR center) or lateral (0-150 bp from HNAR center) footprints. Using the octiles defined above (O1 being the most open, O8 most closed at dome stage), we centered the genomic regions of each octile on [nucmax], oriented as in Fig. 3H, and compared nucleosome prediction values, PT°, MNase signal, and localization of specific motifs between the octiles (Fig. 5A-D). Nucleosome prediction values and PT decreased from O1 to O8 (Fig. 5A,B). The abundance of *nanog, sox*, *C2H2,* and *bHLH* motifs, localized within the medial nucleosome footprint, did not significantly change among octiles (Fig. S19). Thus, specific TF binding within the medial footprint displaces nucleosomes laterally, but it is not sufficient to deplete them (as in octiles O6-O8, Fig. 5C). In contrast, *pou:sox* motif on the lateral footprint was enriched in open octiles (O1-O3, Fig. 5D,E), suggesting that this motif in +90 bp position is critical for nucleosome depletion. Post-ZGA, Pou5f3 bound at +90 bp (Fig. 5F), while Nanog and ATAC-seq peaks were localized more centrally (Fig. 5G,H). We concluded that post-ZGA nucleosome depletion involves two overlapping nucleosome footprints and is therefore distinct from pure nucleosome-mediated cooperativity mechanism. Specific Nanog and Pou5f3 binding to the medial and lateral footprints, respectively, blocks nucleosome assembly on both nucleosome positioning sequences and leads to nucleosome depletion.

**Figure 5.**
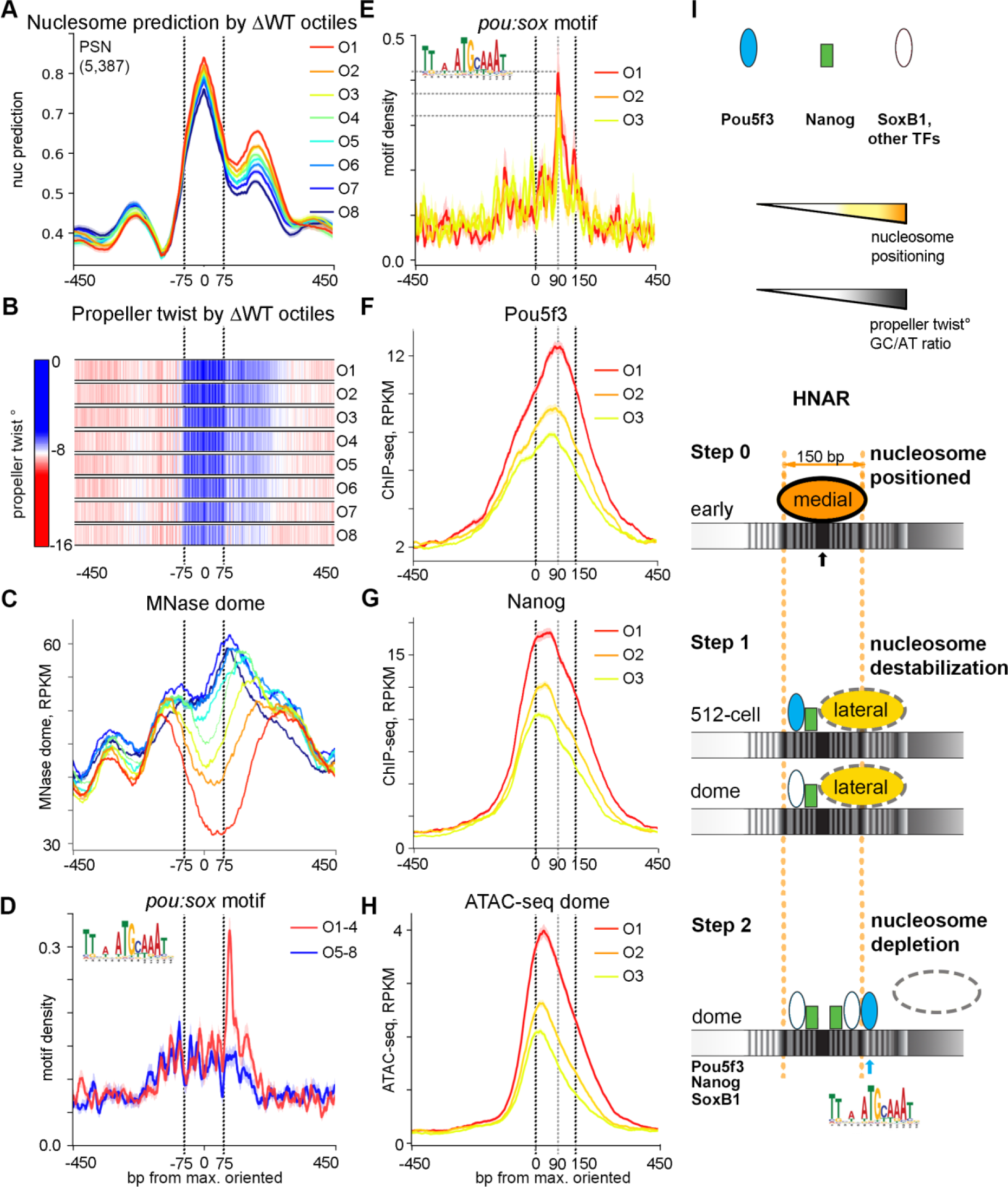
Post-ZGA nucleosome depletion requires Nanog central binding and Pou5f3 binding to +90 bp position on HNAR. PSN group regions ranked into octiles as in Fig. 2, aligned on [nucmax] and oriented as in Fig. 3H. Black dotted lines mark the borders of the medial nucleosome footprint in (*A-D*) and of the lateral nucleosome footprint in (*E-H*). (*A*) Nucleosome prediction profiles per octile. (*B*) Propeller twist heat maps per octile. (*C*) MNase (dome) per octile; note nucleosome displacement to the lateral footprint in O6-O8. (*D,E*) *Pou:sox* motif density, bp motif per bp sequence. (*D*) *Pou:sox* motif peak at + 90 bp is present in open O1-O4 but not in O5-O8 octiles. (*E*) *Pou:sox* motif abundance at +90 bp position decreases O1>O2>O3. (*F)* Pou5f3 binds at the +90 bp position in octiles O1-O3. (*G*) Nanog binds centrally in O1-O3. (*H*) ATAC-seq in O1-O3. (*I*) Two-step nucleosome destabilization-depletion model (see main text). Black and blue arrows show 0 (center) and +90 bp position Pou5f3 binding on HNAR.

### Two-step nucleosome destabilization-depletion model

We summarize our findings in a two-step HNAR-centered nucleosome destabilization-depletion model (Fig. 5I). Initially, in the early pre-ZGA cell cycles, nucleosomes assemble on the strongest medial nucleosome positioning sequence of HNAR (Fig. 5I, Step 0). At 512-cell stage, Pou5f3, Nanog, and possibly other non-histone DNA binding proteins non-specifically destabilize nucleosomes on central nucleosome footprints of HNARs and shift them laterally (Fig. 5I, Step 1, 512-cell). At dome stage, nucleosomes are destabilized by Nanog but not by Pou5f3 (Fig. 5I, Step 1, dome). In the second step, which occurs only after ZGA, simultaneous specific Nanog binding within the medial and Pou5f3 binding within the lateral nucleosome positioning sequence deplete nucleosomes from HNAR (Fig. 5I Step 2, dome).

### HNAR model is applicable to the mammalian system

We tested if the HNAR model is applicable to the mammalian homologs Pou5f3 and Nanog. Experiments using POU5F1, SOX2, KLF and MYC (OSKM mix, Takahashi and Yamanaka 2006) for reprogramming of human or mouse fibroblasts to induced pluripotent stem (iPS) cells suggested different modes of human POU5F1 and mouse Pou5f1 interactions with chromatin. Soufi et al (2015) showed that at the initial stage of human fibroblasts reprogramming POU5F1 targets its partial motifs directly on nucleosomal DNA. Chronis et al. (2017), on the other hand, did not find evidence for direct nucleosome targeting by Pou5f1 in the reprogramming experiment in mouse cells. We addressed if the differences between species can be explained by sequence composition of POU5F1/Pou5f1/Pou5f3 TFBS and location of their motifs on HNARs. We recovered human partial nucleosomal motif (Soufi et al., 2015) and canonical *pou:sox* motifs using mouse and human ES cell ChIP-seq data for POU5F1 and Pou5f1 (Kunarso et al. 2010, Whyte et al. 2013, see Fig. 6A for logos, Table S2 for matrices). We then compared the relative abundances of human partial motif (P), mouse canonical (M) and human canonical (H) motifs across POU5F1/Pou5f1/Pou5f3 ChIP-seq data sets from human and mouse ES cells (Kunarso et al. 2010,Whyte et al. 2013), reprogramming experiments (Soufi et al. 2012, Chronis et al. 2017), and zebrafish embryos. The results indicated that only human POU5F1 can bind the partial nucleosomal motif while canonical motifs are recognized by all Pou5f1 homologs (Fig. 6B-D).

**Figure 6.**
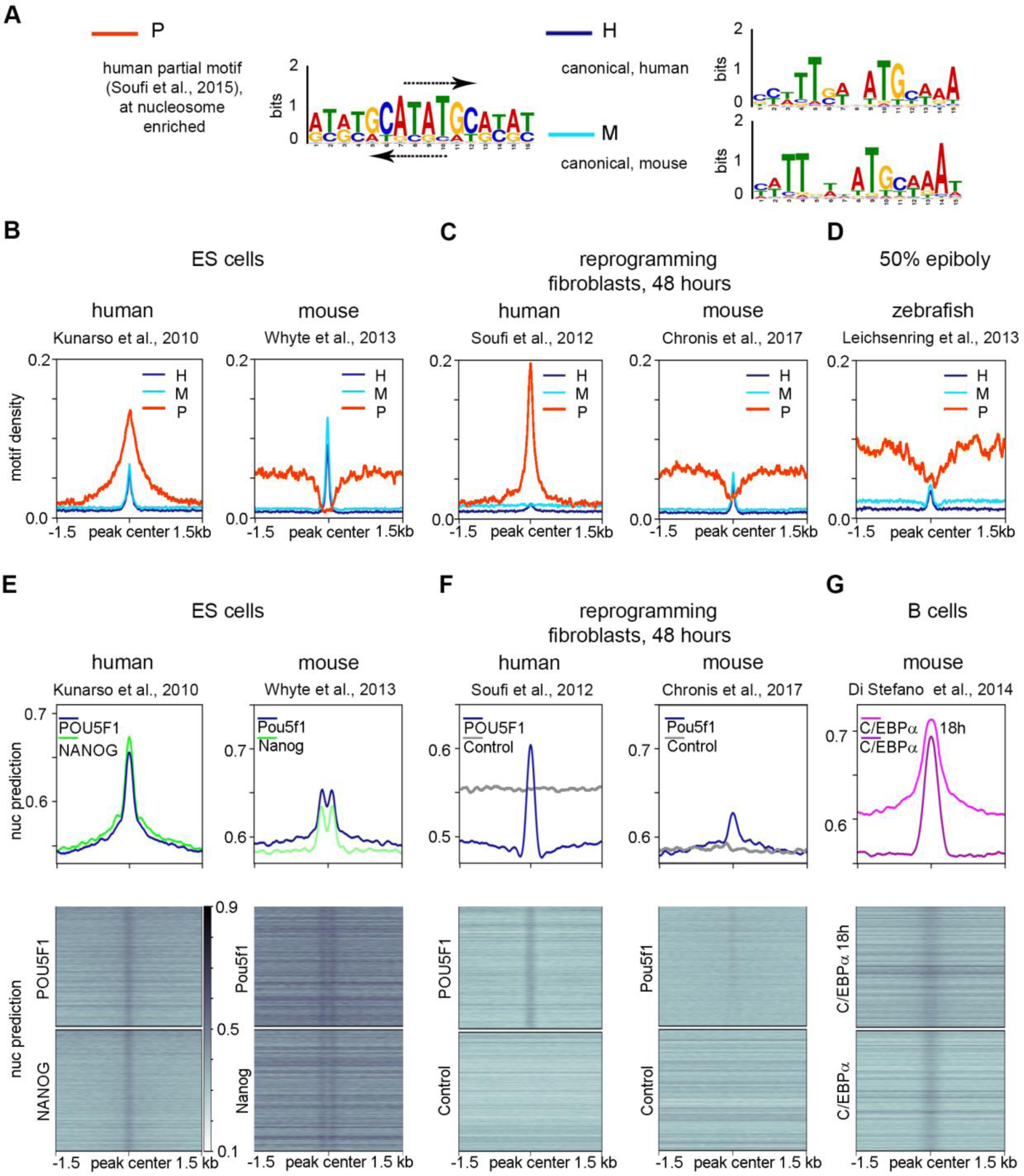
Species-specific chromatin recognition features of POU5F1 homologs. (*A*) Logos of motifs recognized by human POU5F1 (P-partial and H-canonical), and by mouse Pou5f1 (M-canonical). Arrows show the matches to the partial nucleosomal motif of Soufi et al. (2015) on the forward and reverse strands of human palindromic motif. (*B-D*) Densities of partial human motif (P), human canonical motif (H) and mouse canonical motif (M) in the POU5F1/Pou5f1/Pou5f3 ChIP-seq data from the sources indicated on top. (*E-G*) Nucleosome predictions for the ChIP-seq genomic regions from the sources indicated on top.

Nucleosome-binding preferences of POU5F1 protein could indicate that it binds to the HNAR center in human, and their absence mouse Pou5f1 could indicate that it binds outside of HNAR center. To test this, we examined the distribution of the predicted nucleosome occupancy around POU5F1/Pou5f1 ChIP-seq data sets (Fig. 6E,F). Single peaks of predicted nucleosome occupancy in human ES cells indicated that POU5F1 is located on the HNAR center, while bimodal peaks in mouse ES cells indicated that Pou5f1 binds HNAR, but it is located at a distance from the HNAR center (Fig. 6E). In reprogramming experiments, POU5F1, but not Pou5f1, could bind HNAR centers in a way similar to ES cells (Fig. 6F). HNAR destabilization-depletion model plausibly explains the differences in pioneer properties of mouse and human Pou5f1 orthologs: POU5F1, but not Pou5f1 specifically binds to the palindromic motif on HNAR center which may destabilize and shift strongly positioned nucleosomes. C/EBPα TF poises mouse B-cells for rapid reprogramming into induced pluripotent stem cells by OSKM (Di Stefano et al. 2014). C/EBPα was strongly bound to HNAR centers in two experimental conditions (Fig. 6G, data from Di Stefano et al. 2016), suggesting that it may destabilize strongly positioned nucleosomes. We concluded that the HNAR model of zebrafish ZGA is applicable to other systems.

## Discussion

We show that the affinity of both the TFs Pou5f3 and Nanog and nucleosomes to the same locations within the genome is supported by the intrinsic DNA features favoring high *in vitro* nucleosome occupancy. We define the regions spanning 600 bp of high predicted nucleosome occupancy/PT/GC content around the central periodic structure as “High Nucleosome Affinity Regions” (HNARs), to distinguish them from strong nucleosome positioning sequences of one nucleosome length (Fig. 3J). We show that Pou5f3 and Nanog binding is the cause, not a consequence of chromatin opening on HNARs, which defines them as pioneer factors. ZGA TFs are involved in two steps of nucleosome displacement: non-specific competition with histones on strong nucleosome footprints before ZGA and maintenance of open chromatin state by synergistic binding to HNARs after ZGA (Fig. 5I). We think that, although the same TFs are involved in both steps, the two chromatin remodeling steps are driven by separate processes. Connection between the two steps can be described as a weak regulatory linkage (*sensu* Gerhart and Kirschner, 2007).

### Step 1: Destabilization of nucleosomes by non-consensus TF binding to periodic sequences on the HNAR centers

At the 9^th^ cell cycle and before ZGA commences Pou5f3 and Nanog bind to HNARs and reduce the nucleosome occupancy within the central nucleosome footprint in a non-specific manner. We find unlikely that Pou5f3 acts by direct binding to nucleosomes, as it does not recognize human nucleosomal motifs (Fig. 6A-D). Our MNase protocol digested the chromatin to about 80% mononucleosomes. This method depletes “fragile” nucleosomes on the regulatory regions (Iwafuchi-Doi et al. 2016; Mieczkowski et al. 2016). Despite of that, we see mixed ATAC-seq and MNase signals on TFBS: this indicates that the step 1 process is not restricted to regulatory regions and is stochastic. Mixed signals presumably come from different cells: in some cells HNARs are nucleosome-free but in most cells, they are closed by nucleosomes. Pre-ZGA, the Pou5f3 ChIP-seq signal is enriched on all HNAR centers (Fig. S17E). A single-cell, single-molecule imaging study of Pou5f1 and Sox2 demonstrated that nonconsensus interactions with chromatin are central to the *in vivo* search for functional binding sites (Chen et al. 2014). These nonconsensus interactions provide a measurable ChIP signal in a population of cells (Chen et al. 2014). We assume that high sequencing depth of pre-ZGA Pou5f3 ChIP-seq (Leichsenring et al., 2013) allows us to detect nonconsensus Pou5f3 binding pre-ZGA. It was previously shown that multiple TFs outcompete nucleosomes from certain genomic locations (Afek et al. 2015). Nonconsensus binding is enhanced at genomic locations with periodic and symmetric DNA sequences, in which nucleotides of different types alternate (Sela and Lukatsky 2011). HNAR centers are symmetric (Fig. 3J) and are underlain by alternating periodic nucleotide stretches (Kaplan et al., 2009), which makes them likely templates for enhanced nonconsensus binding of multiple TFs. The theoretical model of TF-nucleosome competition on a single nucleosome footprint was suggested by Mirny (2010). One non-trivial prediction of his model is that stabilization of nucleosomes by strong nucleosome-positioning sequence will increase the nucleosome displacement effect of multiple cooperating TFs. This is exactly what we observed: Pou5f3 and Nanog pre-ZGA effects increase with nucleosome prediction strength, and so does ATAC-signal (Fig. 4). Given a non-specific nature of the process, there is no reason to assume that Pou5f3 or Nanog are indispensable for it: other TFs may be also involved. In summary, the simplest explanation for pre-ZGA nucleosome destabilization by Pou5f3 and Nanog is nonconsensus binding to periodic sequences on HNAR centers, where they compete with histones thus preventing nucleosome formation.

### Step 2: Post-ZGA nucleosome depletion by synergistic action of Pou5f3 and Nanog at the presence of SoxB1

We detected widespread alignment of nucleosomes on their DNA-encoded nucleosome positioning signals from pre-to post-ZGA (Fig. 4C, black arrows). Increase of nucleosome positioning strength was previously documented for promoters: zygotic TSS selection grammar is characterized by nucleosome positioning signals, precisely aligned with the first downstream (+1) nucleosome (Haberle et al. 2014). Strengthening of nucleosome attachment to their DNA sites over ZGA is independent of transcription (Haberle et al. 2014, Zhang et al., 2014), but may depend on the elongation of the cell cycle, or on the rapid exchange of maternal nucleosome-associated proteins, such as of conserved maternal linker histones B4/H1M to their zygotic variants (Muller et al. 2002). We speculate that after an initial short phase during the 9-10^th^ cell cycle when histones may be non-specifically outcompeted with TFs and *vice versa* (Joseph et al. 2017), temporally “open” chromatin sites are closed again, unless stabilized by PSN-specific binding (step 2 in our model). Stabilization occurs on HNARs, which are bound by all three TFs (PSN group) and, in addition, carry specific *pou:sox* motif outside of the central nucleosome footprint (+90 bp position at Fig. 5I).

### HNAR model and working hypotheses

The length of HNARs well matches the average size of developmental enhancers (Levine and Tjian 2003). It has been suggested that strong nucleosome positioning signals overlap with the enhancers (Tillo et al. 2010) and pioneer TFs may recognize these signals (Barozzi et al. 2014; Sun et al. 2015). However, it is currently unclear how ubiquitously occurring strong nucleosome positioning signals differ from those contributing to the enhancers. Dinucleotide repeats which we detected on all TFBS (Fig. S2) and which had been previously characterized as general enhancer features (Yanez-Cuna et al., 2014) contribute to the periodic nucleosome positioning signal underlying HNAR center (Kaplan et al., 2009), and at the same time serve as a template for nonconsensus recognition by multiple TFs (Sela and Lukatsky 2011). We hypothesize that most of the TFs present in the cell transiently bind to HNAR centers by default. Further TF-specific DNA shape cues may be decoded as differential specific binding positions of the TFs within the HNAR relatively to its center. We suggest two testable hypotheses:

1) If TF destabilizes nucleosomes in a manner dependent on nucleosome prediction strength of the underlying sequence, this TF is involved in non-specific nucleosome-mediated cooperativity on HNAR center (as in step 1, Fig. 5I).
2) Functional TF-binding motifs involved in transcriptional regulation reside in HNAR outside of medial nucleosome footprint (as *pou:sox* motif in step 2, Fig. 5I).

The 300 bp propeller twist periodic regions, similar in shape, can be recovered from zebrafish, yeast, and human genomes (Fig. 3F). HNARs can be distinguished from random regions by elevated propeller twist and GC/AT ratio (Fig. 3I). These features promote nucleosome assembly and non-consensus binding of various transcription factors: thus, HNAR provides a platform for competition between histones and TFs. In a way, HNARs can be viewed as naturally occurring enhancer templates: adding specific TF binding motifs to HNAR will make an enhancer. We envision, that HNAR concept will be instrumental for future studies of genome regulatory biology of eukaryotes.

## Methods

Materials and methods are in the Supplementary Material.

## Data access

MNase-seq raw and processed data have been submitted to the NCBI Gene Expression Omnibus (GEO; http://www.ncbi.nlm.nih.gov/geo/) under accession number GSE109410. Other processed data is available in the main text or the Supplemental Materials.

## Acknowledgments

We are grateful to Nadine Vastenhouw for sharing MNase-seq protocol, to Wolfgang Driever, Sebastian Arnold, Patrick Lemaire, Meijiang Gao and Marco Ell for commenting the manuscript, to Sabine Goetter for excellent fish care and to Andrea Buderer for administrative support. We are especially grateful to Remo Rohs for maintaining the community access to TFBShape server. This work was supported by DFG-ON86/4-1, DFG-ON86/4-2 and DFG-EXC294 A7-2 for D.O. The Freiburg Galaxy Team is funded by DFG grant SFB 992/1 2012 and BMBF grant 031 A538A RBC.

## Author contributions

D.O. conceived the project, M.V. performed all the experiments, D.O, L.Y and B.G performed the data analysis, D.O. and M.V. wrote the manuscript. All authors edited the manuscript.

