## Supplemental Material for "Pou5f3, SoxB1 and Nanog remodel chromatin on High Nucleosome Affinity Regions at Zygotic Genome Activation"

**This PDF file includes:**

Materials and Methods
Supplementary Text
Figures S1 to S19
Tables S3 to S10
Captions for Data Tables S1 and S2

**Other Supplementary Materials for this manuscript include the following:**

Data Tables S1 and S2

### Supplementary Methods

#### Zebrafish maintenance and embryo collection

Wild-type fish of AB/TL strain were raised, maintained and crossed under standard conditions as described by Westerfield (Westerfield 2000). The mutant *spg*$^{m793}$ and *nanog*$^{m1435}$ lines were maintained as described previously (Lunde et al. 2004; Veil et al. 2018). Embryos obtained from crosses were collected within 10 minutes and raised in egg water at 28.5°C. Staging was performed following the 1995 Kimmel staging series (Kimmel et al. 1995). Stages of MZ*spg*$^{m793}$ and MZ*nanog*$^{m1435}$ embryos were indirectly determined by observation of wild-type embryos born at the same time and incubated under identical conditions. All experiments were performed in accordance with German Animal Protection Law (TierSchG).

#### MNase digestion and sequencing

Around 200-400 wild-type, MZ*spg* and MZ*nanog* embryos were staged to 512-cell (2.75 hours post-fertilization) or dome (4.3 hours post-fertilization) stage. Embryo fixation and the MNase digestion was performed as described (Zhang et al. 2014). The yield of and degree of digestion was controlled using Agilent High Sensitivity DNA Kit on Agilent Bioanalyzer, according to manufacturer instructions. Chromatin was digested so that it contained 80% mononucleosomes (Supplemental Fig. S4). Libraries were prepared using the Illumina sequencing library preparation protocol and single-end sequenced on an Illumina HiSeq 2500 by Eurofins Company (Germany).

#### Library preparation and sequencing

Libraries were prepared using the Illumina sequencing library preparation protocol and single-end sequenced on an Illumina HiSeq 2500 by Eurofins company (Germany).

#### Mapping

All sequenced reads were mapped back to the zebrafish Zv9 assembly using Bowtie2 (Langmead and Salzberg 2012) on Galaxy server (Afgan et al. 2016) in Freiburg (https://galaxy.uni-freiburg.de). Numbers of mapped reads are listed in Table S3. BAM files were converted to BED format using BEDTools (Quinlan and Hall 2010 wrapped into the Galaxy by Bjoern A. Gruening (2014) Galaxy wrapper (https://github.com/bgruening/galaxytools)). To create the files for scoring and visualization of the nucleosome profiles from single end sequencing data, we used the strategy previously described by Zhang et al. (2008). All mapped reads were extended to 147 bp in their 3' direction and truncated to the middle 61 bp. Original mapping BAM files (Zv9) and processed Bigwig files (Zv9 and GRCz11) are available under GEO accession number GSE109410.

#### Selection of TFBS and control groups for analysis

Lists of 6,670 post-ZGA Pou5f3, 7,747 pre-ZGA Pou5f3 and 5,924 SoxB1 ChIP-seq peak genomic coordinates were taken from Leichsenring et al. 2013. Mapped ChIP-seq reads for Pou5f3 pre- and post-ZGA and SoxB1 post-ZGA were from GSE39780 series in GEO NCBI**.** 14,010 dome stage Nanog ChIP-seq peak coordinates in Zv8 genome assembly were taken from Xu et al. 2012, and converted into 13,775 Zv9 peaks using LiftOver utility from the UC Santa Cruz Genome Browser. Raw Nanog ChIP-seq data were uploaded from GSE34683 series in GEO NCBI and mapped to Zv9 genome assembly. To create the group of control genomic regions, we first calculated the genomic distances from 7,500 randomly taken Pou5f3, SoxB1 and Nanog peaks to the transcription start site (TSS) of the closest ENSEMBL transcript and then took the genomic regions at the same distances from randomly picked ENSEMBL transcripts. Previous analysis of the zebrafish promoters demonstrated the genome-wide formation of prominent nucleosome-free regions upstream of the promoters at ZGA (Haberle et al. 2014; Zhang et al. 2014). To exclude the overlap with these nucleosome-free regions we removed the peaks +/-1 kb from the annotated promoters of ENSEMBL transcripts. ChIP-seq and MNase-seq data sets were uploaded to seqMINER (Ye et al. 2011), which allows to simultaneously visualize and score nucleosome and ChIP-seq signal distribution around the set of genomic regions. To reduce the heterogeneity of the analyzed ChIP-seq lists, we processed them in a standard way described below. First, we removed the ChIP-seq peaks, which were falling to the regions of very high MNase-seq occupancy (top 1% of the reads/bp in each MNase sample within any 20 bp in +/- 1 kb from the peak center), with the purpose to exclude the heterochromatin and repetitive regions. To assign the single (P, S, N), double (PS, SN, PN) or triple (PSN) occupancy to the post-ZGA TFBS, we used ChIP-seq BAM files to calculate TF occupancy for P, S and N separately in the central 320 bp (mean peak width) and average from two 320 bp flanks (background). For each genomic region in TFBS list, the region was scored as negative for TF binding, if peak/background signal ratio was less or equal to the arbitrary cutoff 1.3, and positive, if the peak/background ratio was more than 1.3. After the filtering, one quarter of post-ZGA TF peaks (6,139 regions) were removed from the original list of 26,369 regions. To the remaining list, 6,248 Pou5f3, 3,301 Sox2 and 2,437 Nanog binding events were assigned as positive in combinations. We did not assign SoxB1-only group, as SoxB1 binding overlapped with some levels of Pou5f3 or Nanog on all regions, due to deliberately low cutoff 1.3 we used. Nanog only bound and Pou5f3 only bound groups (N and P) were significantly large in spite of the low cutoff 1.3. To assign Ppre (pre-ZGA Pou5f3 – only) group, overlaps between Pou5f3 pre- with any of post-ZGA TFBS were removed, leaving 5,170 genomic regions. Genomic coordinates of the resulting 8 groups of genomic regions (PSN, PS, PN, SN, P, N, Ppre and control) are listed in the Table S1 and were used for further analysis.

**Motif finding and analysis**

To find all the motifs specifically enriched in our data, we used MEME suite at http://meme-suite.org (Bailey et al. 2009). We sorted the TFBS by descending ChIP-seq signal for each TF separately. Top 1,000 60 bp wide sequences and top 270 220 bp wide sequences for post-ZGA Pou5f3, SoxB1 and Nanog, and for Pou5f3 pre-ZGA were used as an input for MEME for de-novo motif finding program (Bailey et al. 2009) with parameters:

| | |
|---|---|
| Motif Site Distribution | ZOOPS: Zero or one site per sequence |
| Maximum Number of Motifs | 10 |
| Minimum Motif Width | 6 |
| Maximum Motif Width | 25 |

with or without enabling palindromic search. All the derived motifs were combined in one list. Motifs which were too similar to the others were removed using MAST (Gupta et al. 2007) with matrix correlation threshold 0.9. The remaining 14 motifs (PWMs listed in Table S2) were compared to JASPAR database using Tomtom (Gupta et al. 2007), to the known Nanog, Pou5f1/Pou5f3, Pou5f1/SoxB1 and SoxB1 motifs (Remenyi et al. 2003; Loh et al. 2006; Chen et al. 2008; Salmon-Divon et al. 2010; Xu et al. 2012; Leichsenring et al. 2013) and enhancer-specific motifs (Yanez-Cuna et al. 2014) and subdivided into 7 groups: *pou:sox* (3 motifs), *nanog1* (1 motif), *nanog2* (5 motifs), *sox* (3 motifs), enhancer-associated dinucleotide repeats (*gaga* repeat and *tgtg* repeat) , *atss* repeat, C2H2 Zn finger motif and bHLH motif (Fig. S2). Genomic coordinates of the individual occurrences of each motif in +/- 1.5 kb from the center of all TFBS regions were obtained using FIMO (Grant et al. 2011) with p-value threshold $10^{-4}$. The genomic coordinates of motifs were saved as a BED file, converted to bedgraph format with BEDTools (Quinlan and Hall 2010), and used for plotting the heat maps in seqMINER (Ye et al. 2011) or deepTools2 ( (Ramirez et al. 2016), Fig. 1B, Fig. S2. The motifs from mouse and human ES cells (Table S2, Fig.6) were found and processed the same way. To score *pou:sox* and *nanog1*&*2* motifs per genomic region, overlapping motifs within *pou:sox* group and *nanog1* and *nanog2* groups, respectively, were merged together and trimmed to 20 bp. The numbers of non-redundant matches to *pou:sox* and *nanog* per each 320 bp long TFBS was scored in Galaxy and listed in Table S1.

**Comparative nucleosome occupancy plots**

Analysis was done on Galaxy instance in Freiburg (now Galaxy europe). 6 BED files with MNase-seq data for two stages and three genotypes were converted to BigWig format using BEDTools. BigWig files of log2 fold change differences between the mutants and wild-type, or the morphants and wild-type, were obtained using bigwig compare in deepTools2. DeepTools2 were used for plotting (Ramirez et al. 2016).

#### Zebrafish data sets from other publications

Processed data were used as supplied by publisher. ATAC-seq data (Liu et al. 2018): processed BigWig files in Zv9 assembly of three replicates per stage were downloaded from GSE101779. Replicates were merged for subsequent analysis. DNA methylation data (Jiang et al. 2013): processed Wig files for egg GSM1133392 and sperm GSM1077593 DNA methylation in Zv9 assembly were converted to BigWig files and plotted on motifs specified in this study (the results are shown on Fig. S9). Placeholder nucleosome locations (Murphy et al. 2018): processed BED files for pre-ZGA H3K4me1 and H2A.Z in Zv10 assembly were downloaded from GSE95030. Genomic coordinates of TFBS used in this study were lifted to Zv10 using LiftOver utility from the UC Santa Cruz Genome Browser and intersected with pre-ZGA H3K4me1 and H2A.Z- positive regions. TFBS was scored as positive, if at least one bp of 320 bp TFBS overlapped with H3K4me1 and H2AZ regions (the results are shown on Fig. S8). Zebrafish promoters (Haberle et al. 2014): classification to four expression groups was based on SOM clusters in Haberle et al (2014): Clusters 0_4, 0_3,1_4 - early zygotic, clusters 1_0, 1_1, 0_0, 0_1 – late zygotic, cluster 6_6 - non-expressed, the rest of the clusters –maternal-zygotic (throughout active). The results are shown on Fig. S6.

#### Human and mouse data sets

Processed data were used as supplied by publisher. Genomic coordinates of POU5F1 and NANOG ChIP-seq peaks in human ES cells (Kunarso et al. 2010) were downloaded from GSE20650 as BED files. Genomic coordinates of Pou5f1 and Nanog ChIP-seq peaks in mouse ES cells (Whyte et al. 2013) were downloaded from GSE44286 as WIG files, peaks were called using MACS2 (Feng et al. 2012). From the list of POU5F1 and Pou5f3 ChIP-seq peaks, 1,000 regions (60 bp around peak center) were randomly taken for motif finding with MEME (Bailey et al. 2009). Genomic coordinates of POU5F1 ChIP-seq in human foreskin fibroblasts 48 hrs post-induction with OSKM (Soufi et al. 2012) were downloaded from GSM896985 as BED file. Genomic coordinates of Pou5f1 ChIP-seq in mouse embryonic fibroblasts 48 hrs post-induction (Chronis et al. 2017) were downloaded from GSM2417130 as BED file. Genomic coordinates of C/EBPα ChIP-seq in mouse B-cells and mouse B-cells 18 hr after C/EBPα transfection (Di Stefano et al. 2016) were downloaded from GSE52373 as BED files. To create the group of control genomic regions, we calculated the genomic distances from 20,000 randomly taken ChIP-seq peaks to the transcription start site (TSS) of the closest ENSEMBL transcript and then took the genomic regions at the same distances from randomly picked ENSEMBL transcripts.

#### Cross-species comparison of motif frequencies in POU5F1, Pou5f1 and Pou5f3 TFBS

20,000 regions were selected randomly from the POU5F3 and Pou5f3 ChIP-seq lists for human and mouse data sets. Pou5f3 post-ZGA TFBS were taken in order presented in Fig.1A, main text: PSN, PS, PN and P. All TFBS were extended to 3 kb around the center and converted to fasta format. All occurrences of human palindromic motif (P), human canonical motif (H) and mouse canonical motif (M) of all TFBS regions were obtained using FIMO (Grant et al. 2011) with p-value threshold $10^{-4}$. The genomic coordinates of motifs were saved as a BED file, converted to bedgraph format with BEDTools (Quinlan and Hall 2010), and used for plotting the profiles in deepTools2 ((Ramirez et al. 2016), Fig. 6B-D).

#### Nucleosome predictions

Nucleosome prediction program from Kaplan et al. 2009 was integrated into the Galaxy platform using the Galaxy tool SDK planemo (https://github.com/galaxyproject/planemo) and following the best practices for Galaxy tool development (http://galaxy-iuc-standards.readthedocs.io/en/latest/best_practices.html). The tool was uploaded into the Galaxy ToolShed (ref. https://www.ncbi.nlm.nih.gov/pubmed/25001293) and is available at the Galaxy instance in Freiburg. The sequences around TFBS and controls were extended to 10 kb to account for the edge effects; the nucleosome prediction for each base in the middle 1-3 kb were taken for analysis.

#### Nucmax and oriented HNAR plots

The maximal and minimal nucleosome prediction values within 320 bp around TFBS and control regions and their genomic positions are listed in Table S1. Using smaller windows around TFBS (45,

75 and 200 bp) for max. search did not change the 300 bp periodic shape around [nucmax], subsequently revealed by PT. To orient the genomic regions aligned on [nucmax] along ascending nucleosome prediction values, we searched for the min. nucleosome prediction at +/- 160 bp around [nucmax]. If the min. prediction was downstream of [nucmax], we reversed the strand from + to -. The strand for oriented plots is listed in Table S1. Nucleosome predictions were converted to BigWig files and used for plotting in DeepTools2 (Ramirez et al. 2016)

**Propeller Twist shape**

Propeller twist values for individual sequences or aligned groups of sequences were calculated on TFBS shape server at http://rohslab.cmb.usc.edu/TFBSshape/ (Yang et al. 2014)

**Data normalization and statistical analysis**

The sequencing coverage between the samples was normalized as RKPM (reads per million reads per one kilobase). Normalized difference between the mutant and wild-type (Δmut) were calculated as Δmut=((rpkm(mut)-rpkm(wt))/(rkpm(mut)+rpkm(wt)); normalized difference between the stages (ΔWTpost-pre) was calculated as ΔWTpost-pre=(rpkm(WTpost)-rpkm(WTpre))/(rkpm(WTpost)+rpkm(WTpre)). Average RPKM values per 320 bp were taken. Data were analyzed using JMP (SAS Institute 2012 version 10) using one-way ANOVA with transcription factor binding group as the factor, followed by Tukey-Kramer test for pair-wise differences with p-value set to 0.01, as indicated in the supplementary figure legends and tables.

**Supplementary figures**

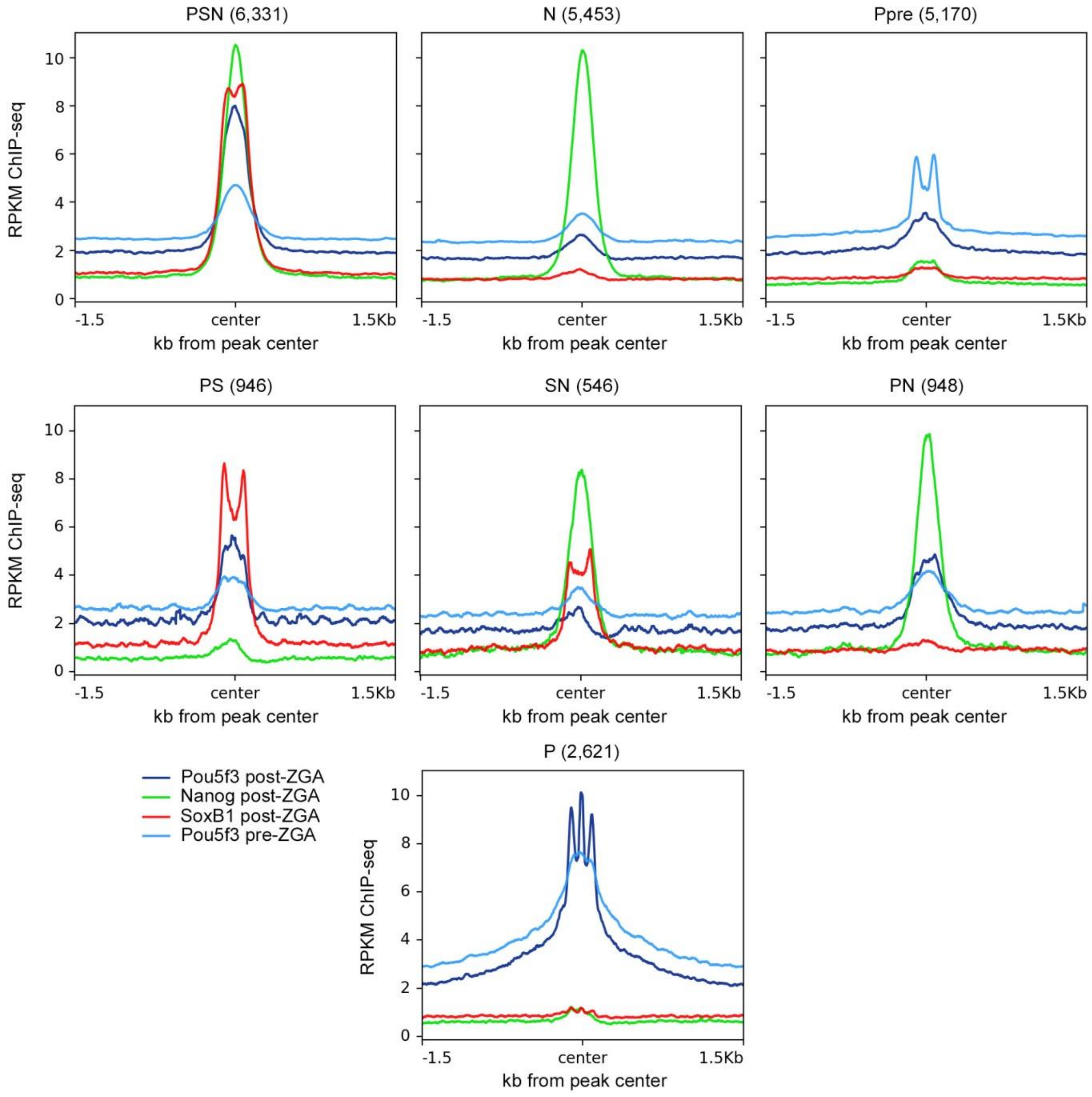

**Fig. S1.**

**Summary TF binding plots for 7 TFBS groups defined in this study.** The graphs show the summary ChIP-seq signal profiles for each TF in all groups. Bin size 10 bp.

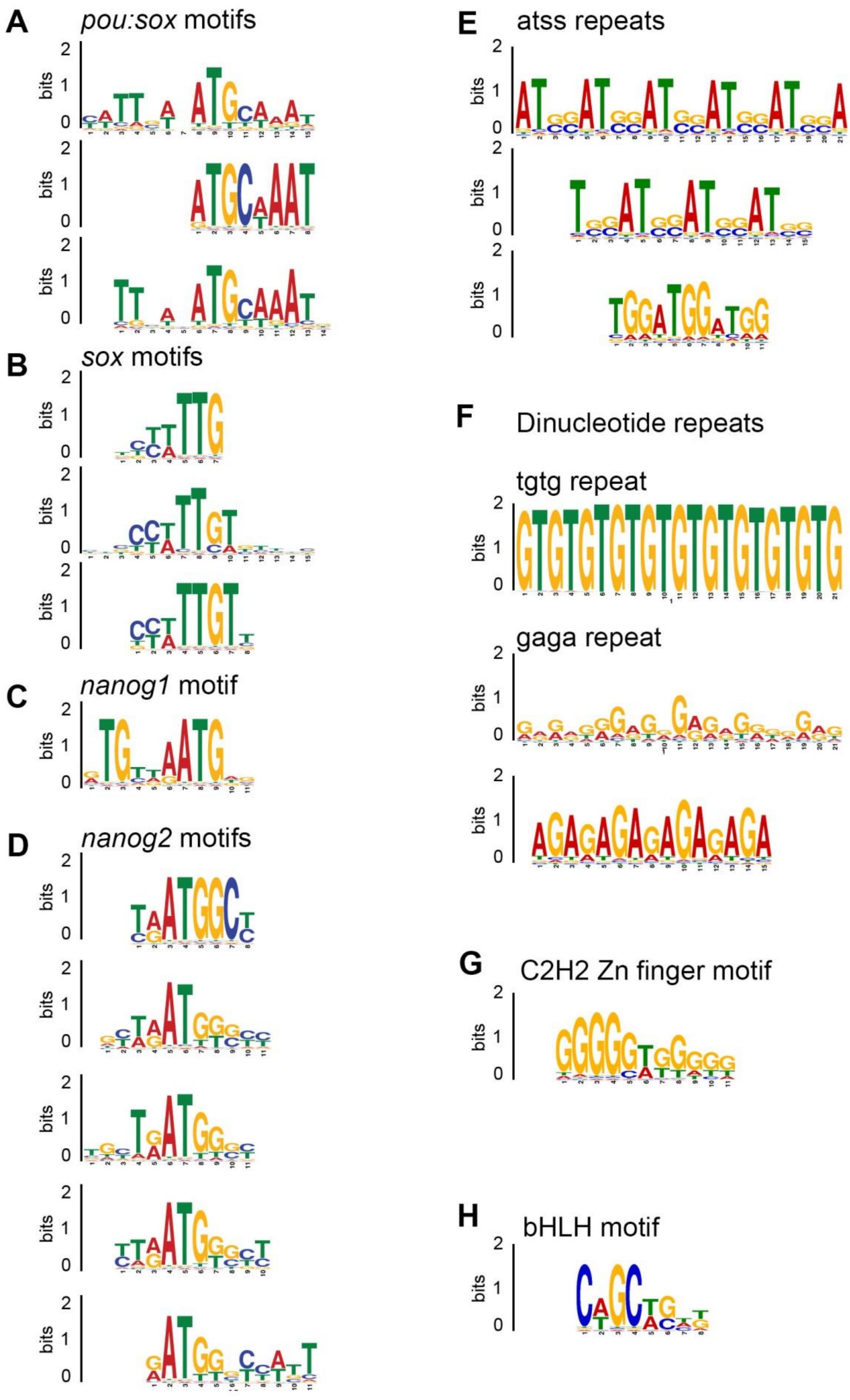

**Fig. S2.**

**Motifs enriched in TFBS from this study.** Sequence logos for specific TF-binding motifs as indicated (see Table S2 for motif matrices).

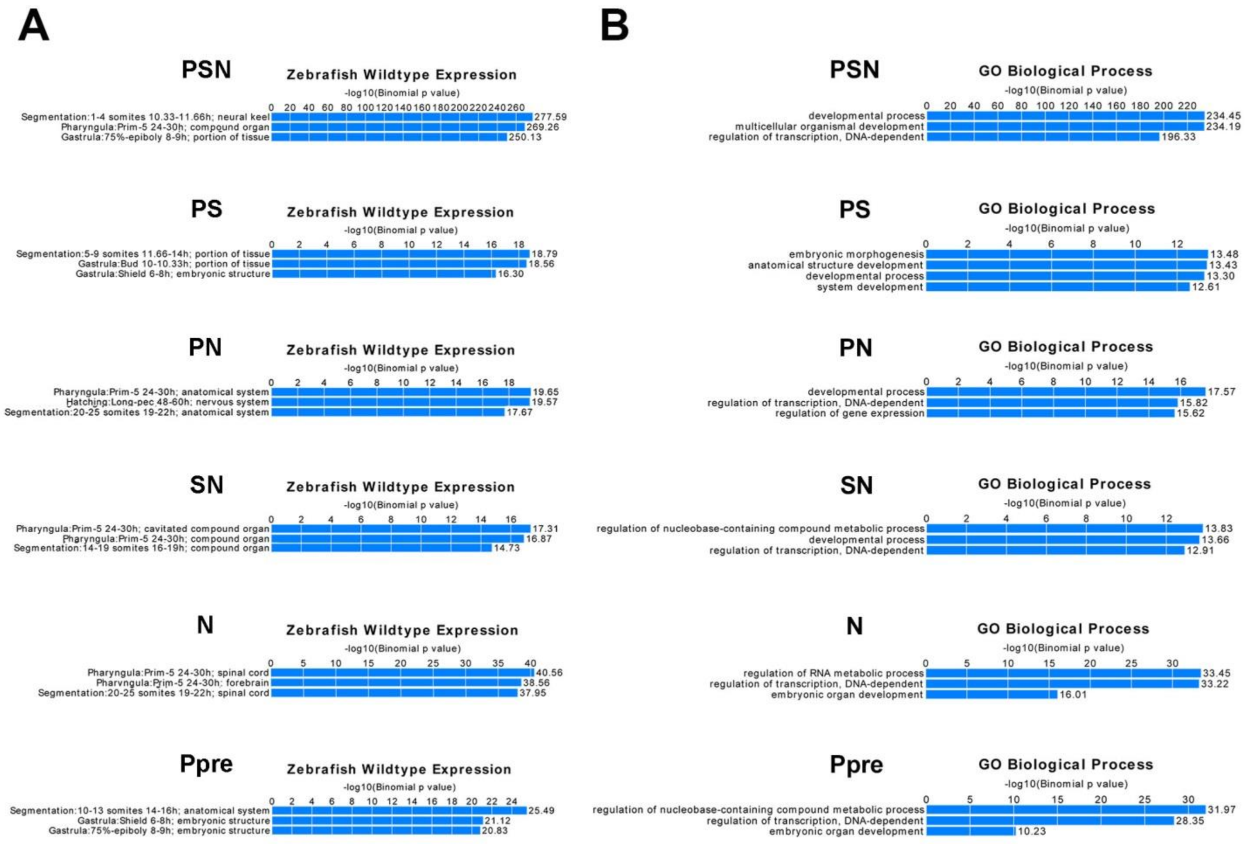

**Fig. S3.**

**6 out of 7 TFBS groups defined in this study are significantly associated with transcriptional regulators and developmental genes.** Selected top categories for Zebrafish WT expression (*A*) and Gene Ontology Biological Process (*B*) for each group from GREAT analysis (McLean et al. 2010). Note that enrichment for PSN group (regions bound by Pou5f3, SoxB1 and Nanog post-ZGA) is the order of magnitude higher than others in both categories. P group of Pou5f3-only post-ZGA binding regions did not show significant enrichments.

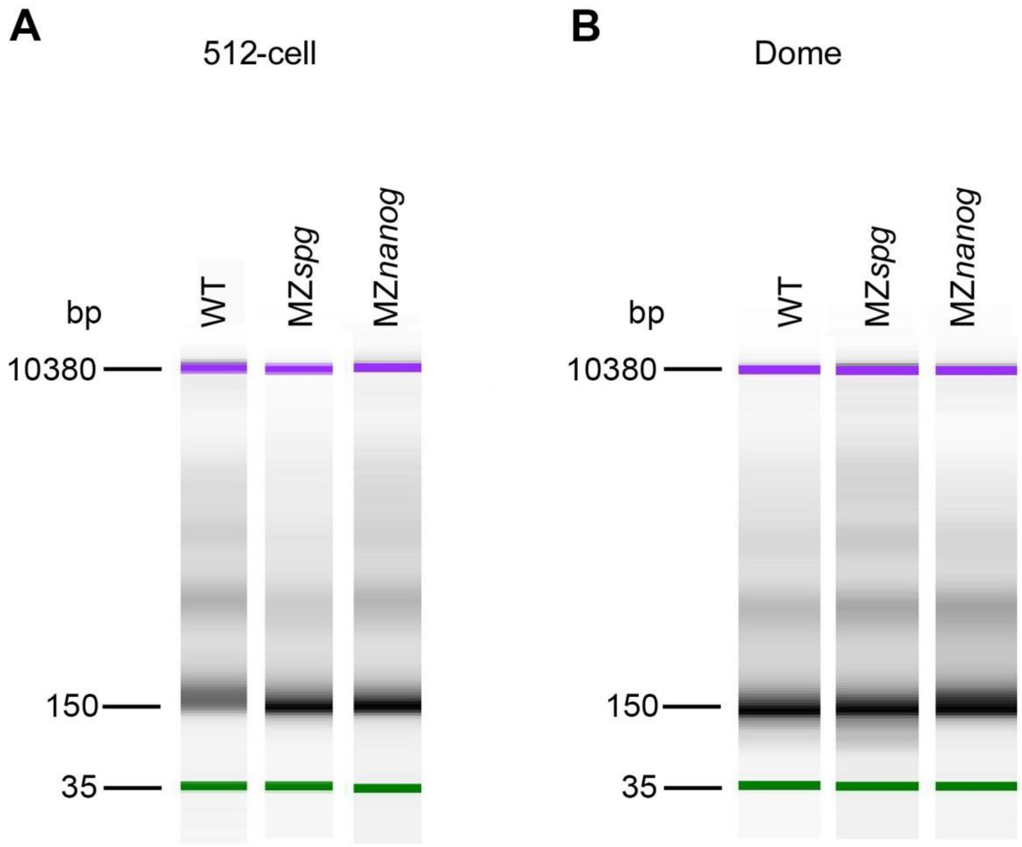

**Fig. S4.**

**Nucleosome fragment length estimation after MNase digestion by using a bioanalyzer (Agilent).** Mononucleosomes are displayed at the size of 150 bp for 512-cell stage (pre-ZGA) (*A*) and dome stage (post-ZGA) (*B*).

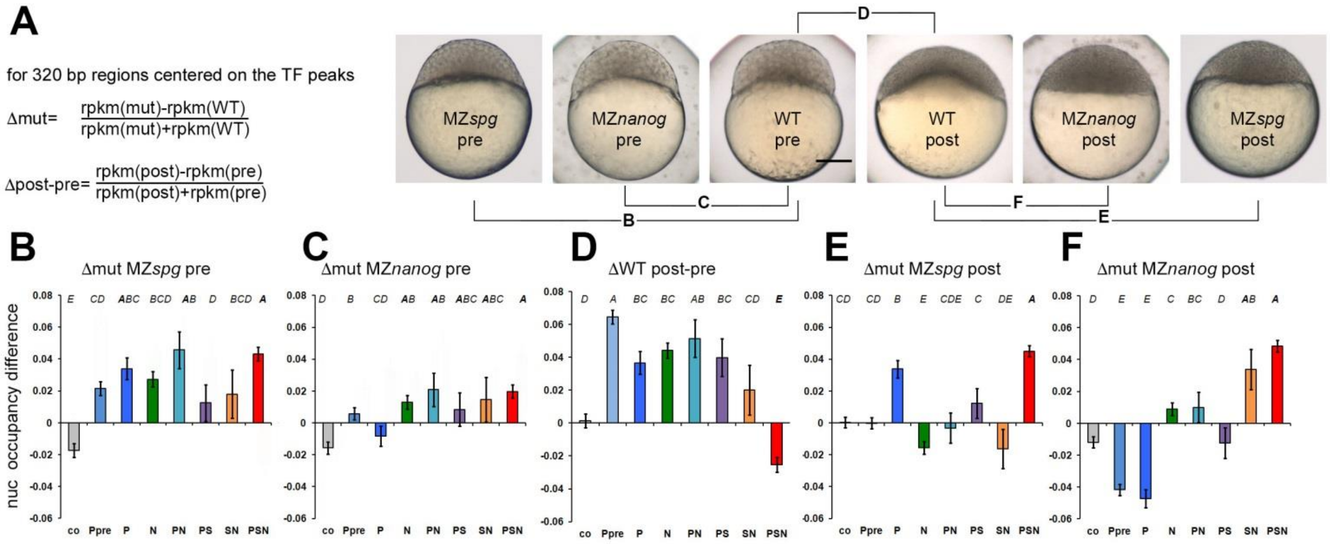

**Fig. S5.**

**Statistics of the nucleosome occupancy change between stages and genotypes.** (*A*-*F*) RPKM values were taken for the 320 bp (mean TF peak width). Error bars are 95% confidence intervals. n(Co) = 6,886, n values for the other groups are indicated in (Fig. 1 main text). Statistical evaluation was done using 1-way ANOVA (Table S4) and Tukey-Kramer test: groups not sharing a letter on the top of the graph are significantly different (P=0.01).

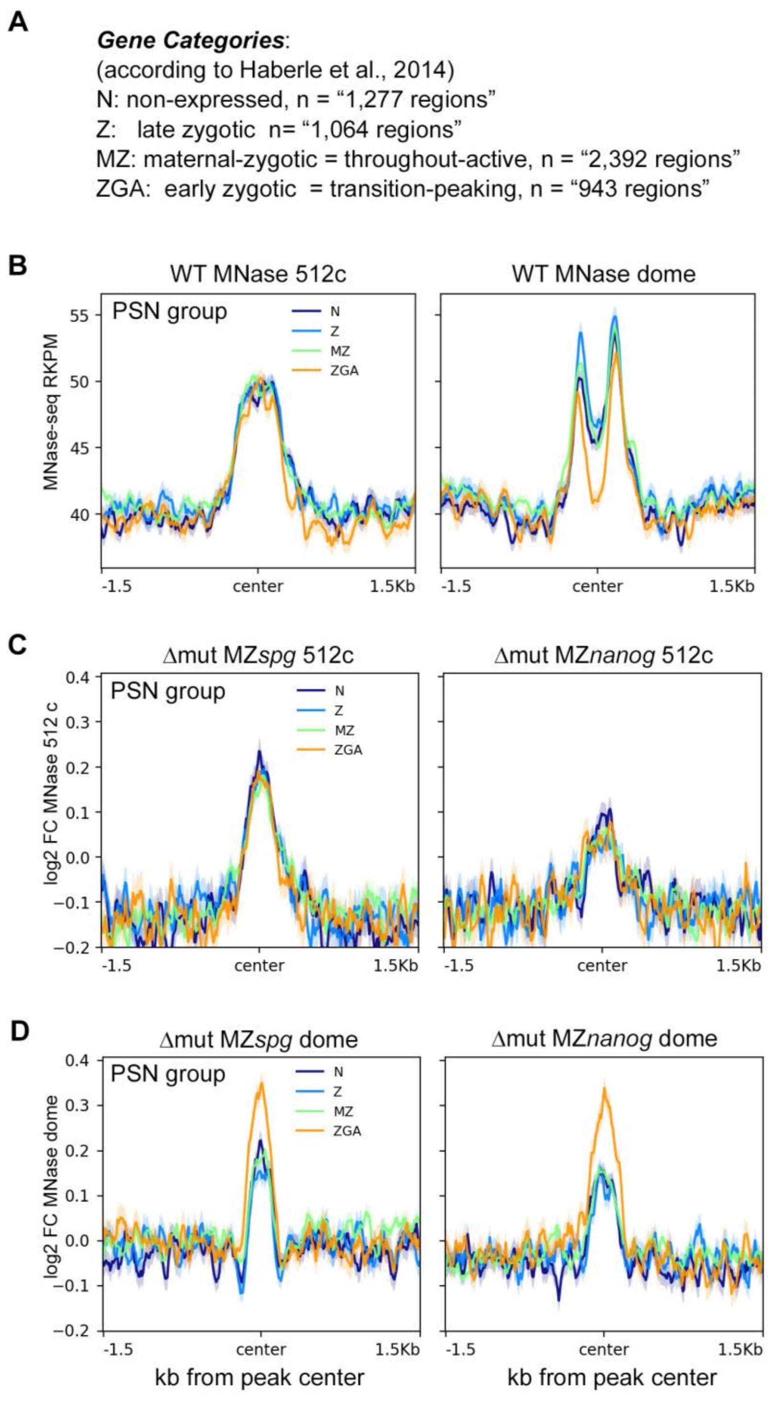

**Fig. S6.**

**Nucleosome displacement by Pou5f3, SoxB1 and Nanog at dome stage occurs preferentially on TFBS which are close to the earliest zygotic genes**. (A) Four expression categories of TFBS by closest promoter *(B)* PSN group, nucleosome occupancy profiles on TFBS center in the wild-type: left – 512-cell stage, right - dome stage. (*C-D*) PSN group, Log2 fold change (FC) between the wild-type and mutant nucleosome occupancies, mutants indicated on top. *(C)* 512-cell stage (*D*) dome stage. No differences by expression group were detected in other TFBS groups (single- or double-TF-occupied regions i.e. N, PS or PN).

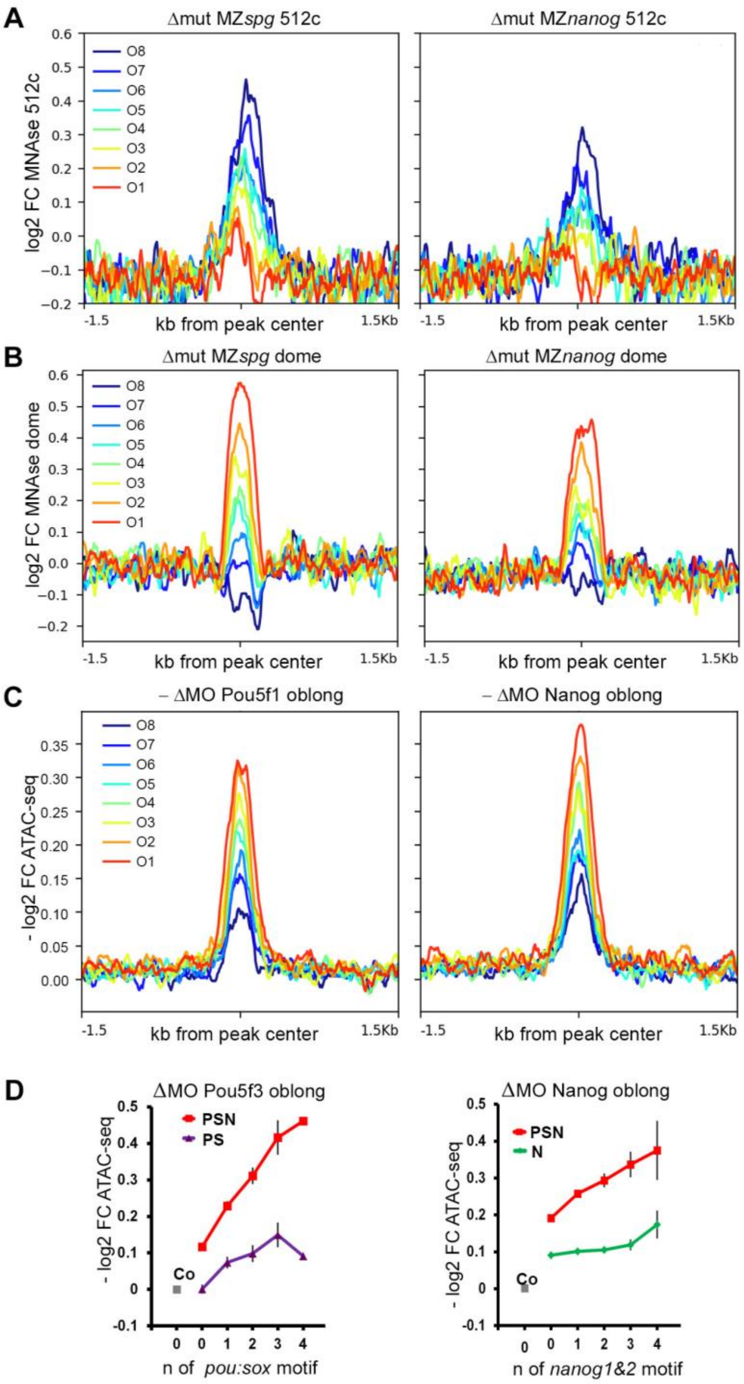

**Fig. S7.**

**Pou5f3 and Nanog effects on nucleosome clearance do not match before and after ZGA. Related to the Fig. 2 of the main text.** (*A*,*B*) Mutant to the wild type differences in MNAse signal per ΔWT octile, PSN group, dome stage**.** Δmut MZ*nanog* and Δmut MZ*spg* are plotted as Log2 fold change between the mutant and wild-type**.** (*A*) 512-cell stage (*B*) dome. (*C)* Wild-type to morphant differences in ATAC-signal per ΔWT octile, PSN group, oblong stage**.** ΔMO Pou5f3 and ΔMO Nanog are plotted as -Log2 fold change (FC) between the morphant and wild-type ATAC-seq signals (Liu et al., 2018) in the PSN regions. Note that ATAC-seq signal is reduced at most in the O1 in both morphants compared to the wild-type. (*D*) ΔMO Pou5f3 and ΔMO Nanog - Log2 fold change between the morphants and wild-type ATAC-seq signals in the genomic regions overlapping 0, 1, 2, 3 and ≥4 motifs and random control (grey dot - Co). FC - fold change.

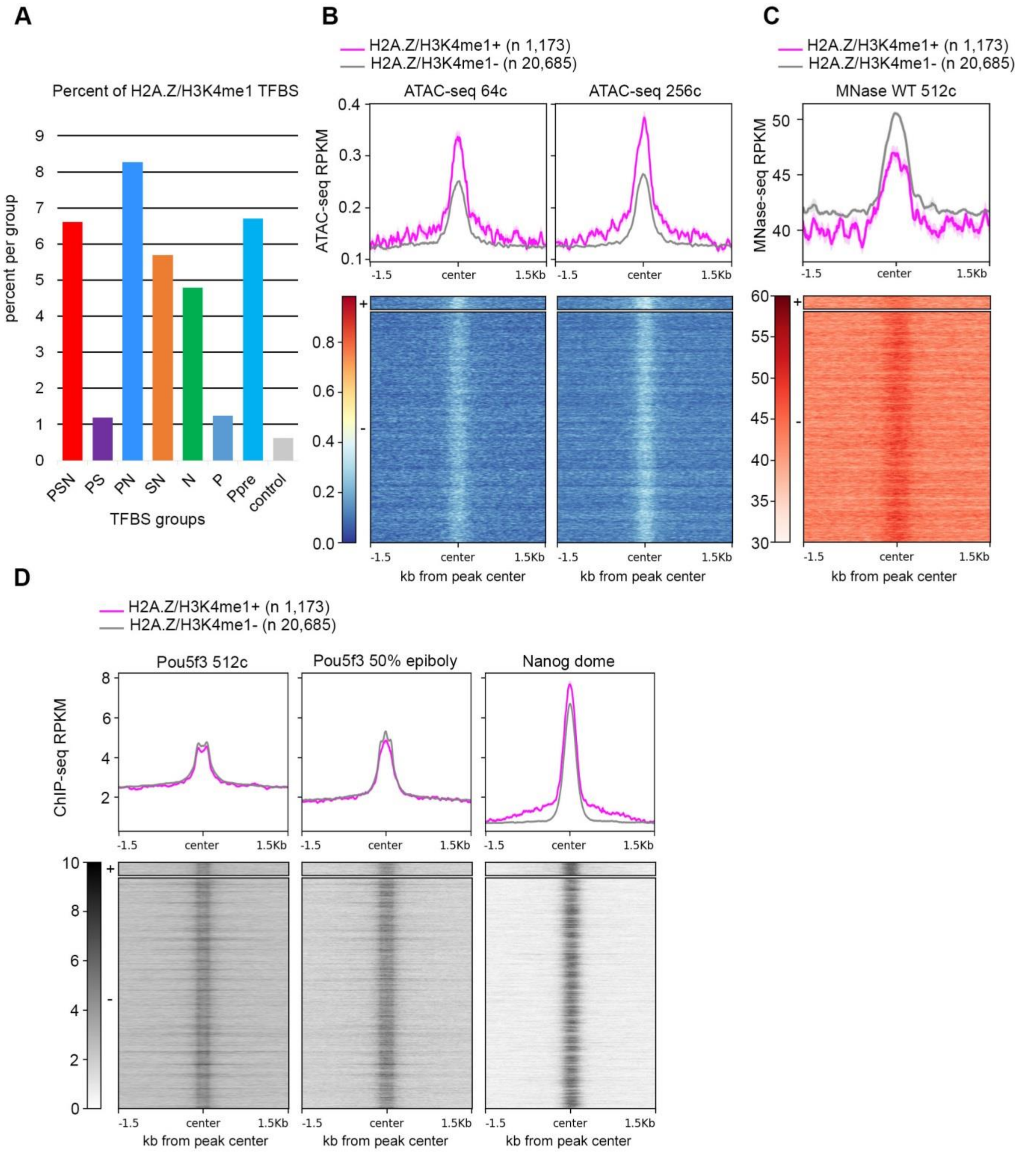

**Fig. S8.**

**Pre-existing H2A.Z/H3K4me1 marks do not influence Pou5f3 and Nanog TF binding.** (*A*) Overlap between TFBS classes and H2A.Z/H3K4me1+ regions per group. (*B)* Pre-ZGA ATAC-seq signal is higher on 5.3% H2A.Z/H3K4me1+ TFBS. (*C*) Pre-ZGA MNase-seq signal is lower on 5.3% H2AZ/H3K4me1+ TFBS. (D) Pou5f3 and Nanog bind H2A.Z/H3K4me1+ and H2A.Z/H3K4me1- TFBS equally.

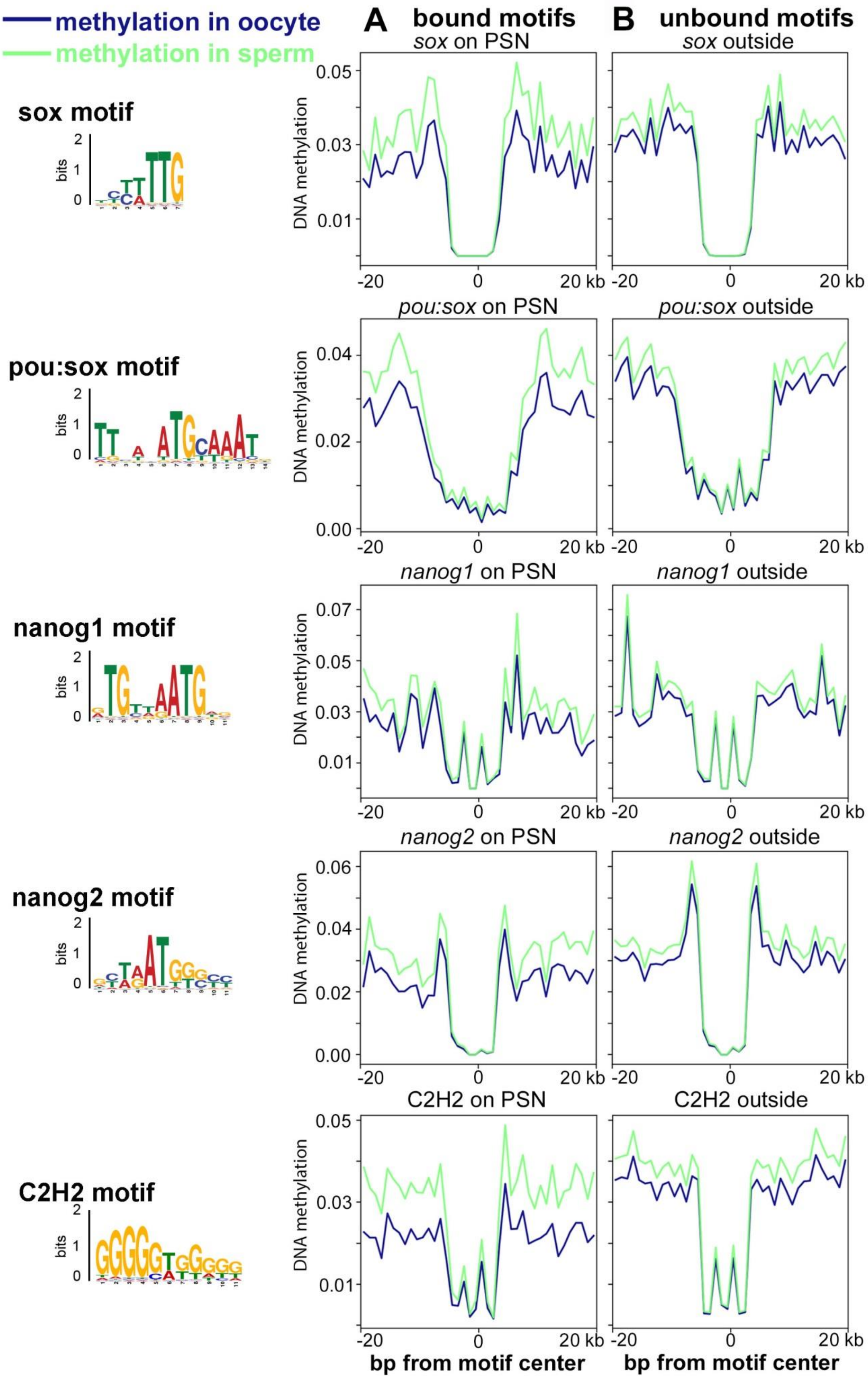
methylation in oocyte
methylation in sperm
A bound motifs
B unbound motifs
sox motif
pou:sox motif
nanog1 motif
nanog2 motif
C2H2 motif
bits
sox on PSN
sox outside
pou:sox on PSN
pou:sox outside
nanog1 on PSN
nanog1 outside
nanog2 on PSN
nanog2 outside
C2H2 on PSN
C2H2 outside
DNA methylation
-20
0
20 kb
bp from motif center

**Fig. S9.**

**Pou5f3 and Nanog bind independently on DNA methylation.** 40 bp genomic regions were centered on the 7-15 bp TF-binding cognate motifs, shown on the left. Mean methylation profiles in the oocyte and in sperm (data from Jiang et al., 2013) are shown in blue and green (top). (*A***)** Methylation profiles of the motifs, bound by TF (within 320 bp of TFBS in the PSN regions). (*B)* Methylation profiles of the random unbound matches for the same motifs (1-1.5 kb from TFBS). No difference in methylation was observed between the sperm and the oocyte, or between bound and unbound motifs.

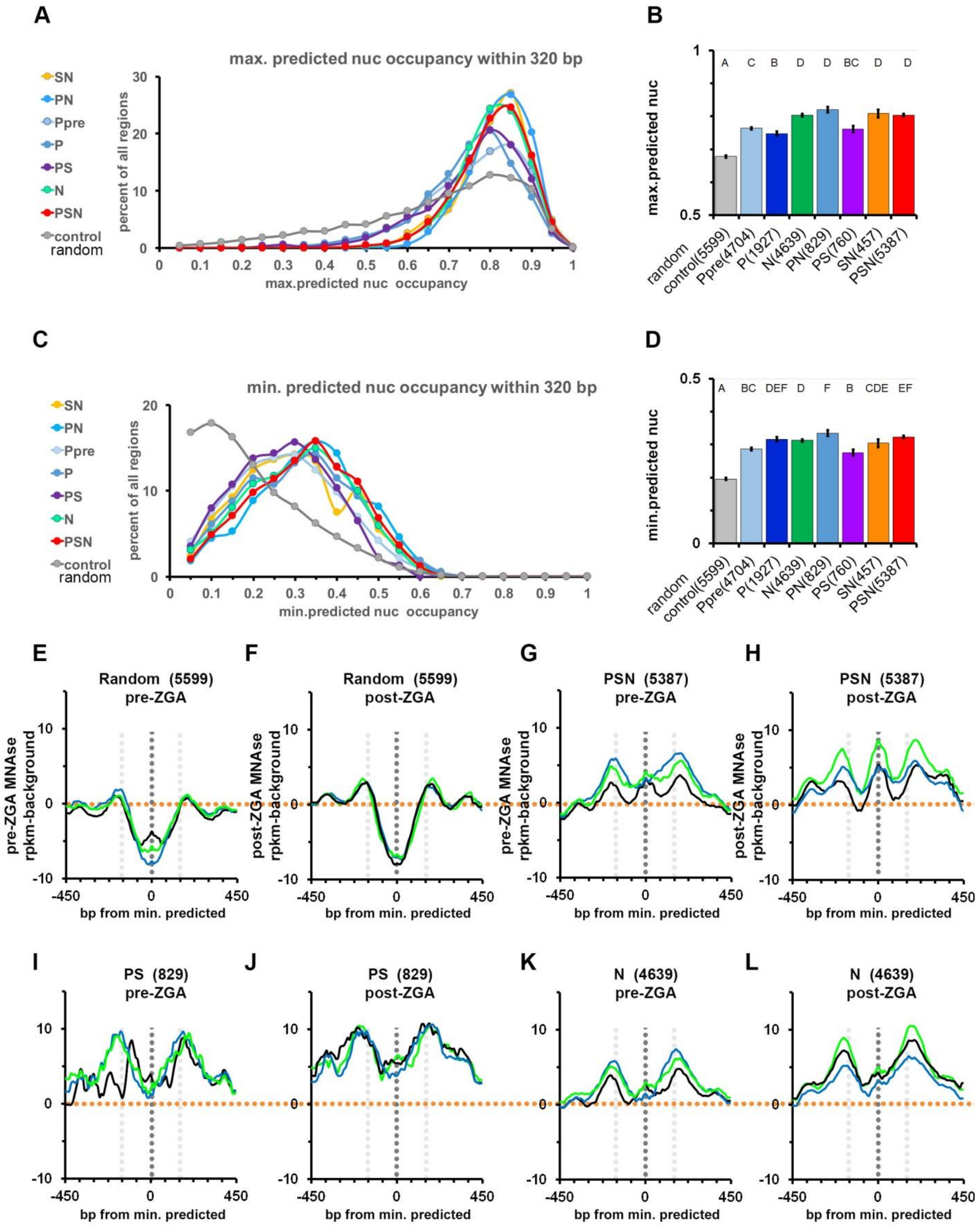

**Fig. S10.**

**Pou5f3, SoxB1 and Nanog bind to the regions of high predicted *in-vitro* nucleosome occupancy which are at least 320 bp long.** (*A-D*) Maximum and minimum predicted nucleosome occupancy value [nucmax] and [nucmin] within each 320 bp long TFBS and control regions were calculated using the program of Kaplan et al. 2009. (*A,C*) The distribution of [nucmax] (*A*) and [nucmin] (*C*) values for 8 groups: Y-axis shows percent of all regions within a group. (*B*,*D*) The differences of [nucmax] (*B*) and [nucmin] (*D*) between the groups (see Tables S7 and S8 for the 1-way ANOVA details). Mean values are shown, error bars are 95% confidence intervals. Letters above the graph represent the results of Tukey-Kramer test: categories not significantly different from each other pair-wise ($p \geq 0.01$) share a letter. (*E*-*L*) Experimental nucleosome occupancy centered on the predicted [nucmin] inter-nucleosomal regions, in the wild-type (black), MZ*spg* (blue) and MZ*nanog* (green) in control (*E*,*F*) and indicated TFBS (*G*-*L*), pre-ZGA(*E*,*G*,*I*,*K*) or post-ZGA (*F*,*H*,*J*,*L*). In control, nucleosomes are absent from [nucmin], as predicted (*E*,*F*); while in PSN, N and PS [nucmin] is occupied by nucleosomes in all genotypes and stages (*G*-*L*). Orange dotted lines show background level, black dotted lines show 0 and gray lines and +/- 150 bp from [nucmin].

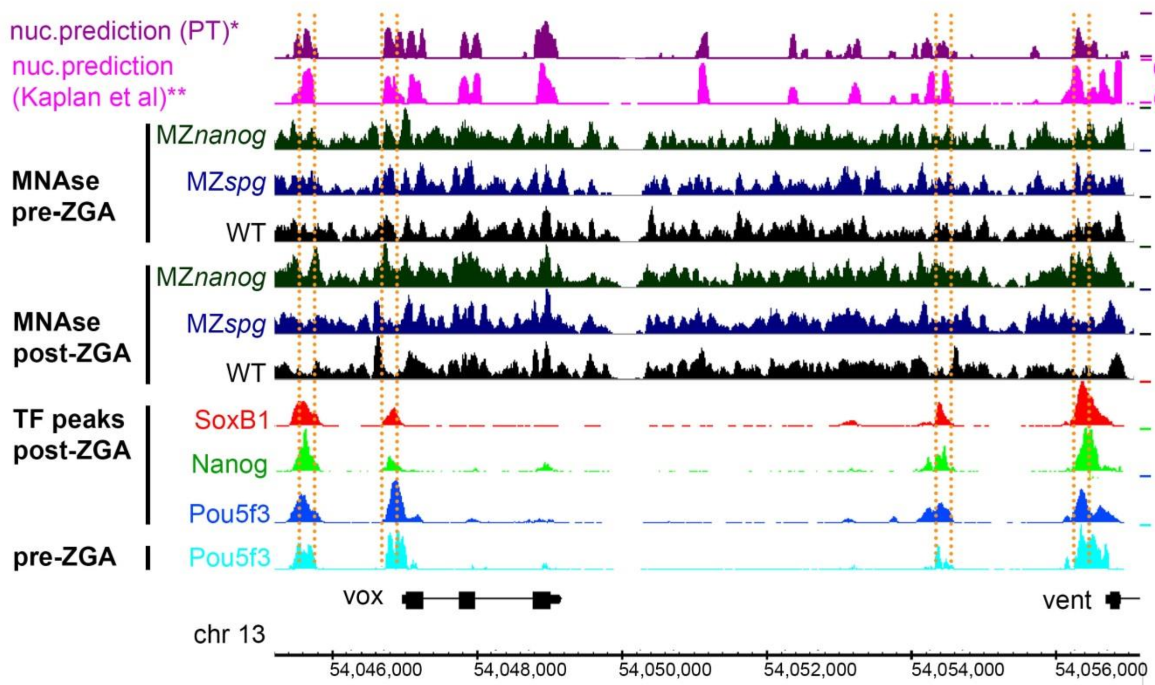

**Fig. S11.**

**Pou5f3, SoxB1 and Nanog bind to the regions of high propeller twist on the *vox* and *vent* enhancers and promoters.** 10 kb genomic region with *vox* and *vent* genes (Zv9). *vox* and *vent* are known transcriptional targets of both Pou5f3 (Reim and Brand 2006; Belting et al. 2011) and Nanog (Perez-Camps et al. 2016; Veil et al. 2018). From top to bottom: PT- propeller twist DNA shape (°), values smoothened with 80 bp moving average, nuc –*in-vitro* nucleosome occupancy prediction (Kaplan et al. 2009), 512-cell, dome stage – pre-ZGA and post-ZGA experimental nucleosome occupancy in the WT (black), MZ*nanog* (green) and MZ*spg* (blue), post-ZGA TF peaks, as indicated, pre-ZGA Pou5f3 TF peaks. Orange lines mark enhancers and promoters of *vox* and *vent*.

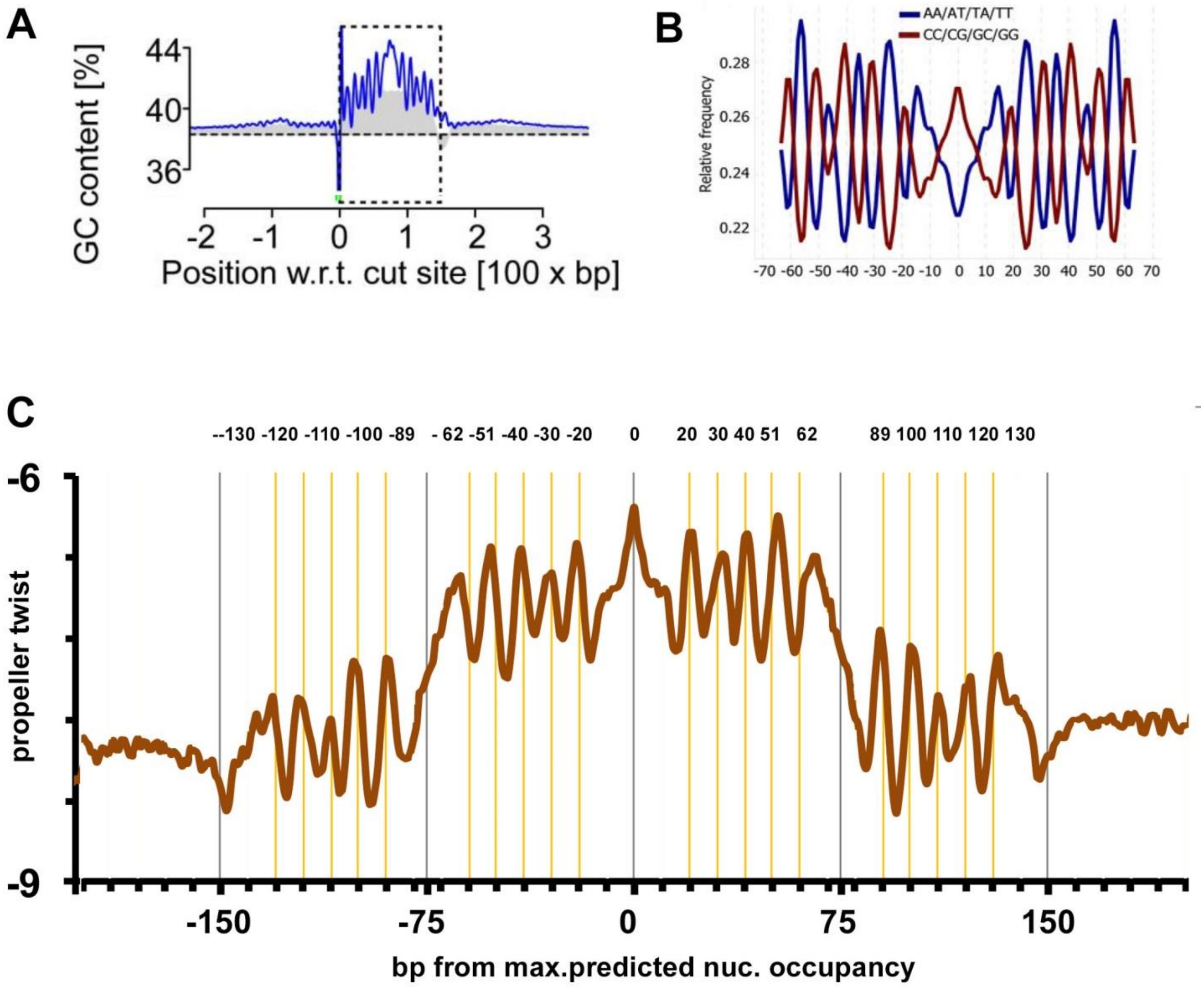

**Fig. S12.**

**Comparison of 10 bp periodicity in 300 bp PT footprint found in this study with the previously characterized GC content periodicity of 150 bp yeast *in-vitro* nucleosome footprints.** (*A*,*B*) Images from (Chung et al. 2010) (*A*) and (Field et al. 2008) (*B*) show the GC content peaks at positions ± 62, ± 51, ± 40, ± 30, ± 20 from the center (dyad) of the nucleosome-bound yeast DNA. These locations correspond to 10 out of 14 positions where the major groove of the DNA faces inwards towards the histone octamer (Chung and Vingron 2009). Images are reproduced under permission of Creative Commons Attribution License. 10 bp periodic oscillations of AT/GC content is a reproducible feature of nucleosomal DNA (Field et al. 2008; Chung and Vingron 2009) reflecting the DNA positioning on the nucleosome core (Satchwell et al. 1986; Richmond and Davey 2003). (*C*) Un-smoothened PT plot of 5,387 PSN group genomic regions, aligned on predicted maximal nucleosome occupancy point [nucmax] (Kaplan et al. 2009). Yellow lines show 10 PT peaks at ± 62, ± 51, ± 40, ± 30, ± 20 which exactly match the AC/GC content fluctuations seen above, and 10 additional PT peaks at ± 130, ± 120, ± 110, ± 100 and ± 89, within the same 10 bp period, which were not described before. PT values were derived using TFBSshape server http://rohslab.cmb.usc.edu/TFBSshape/ (Yang et al. 2014). Average values per 1 bp were plotted. Minor tick marks in the X-axis are 10 bp apart.

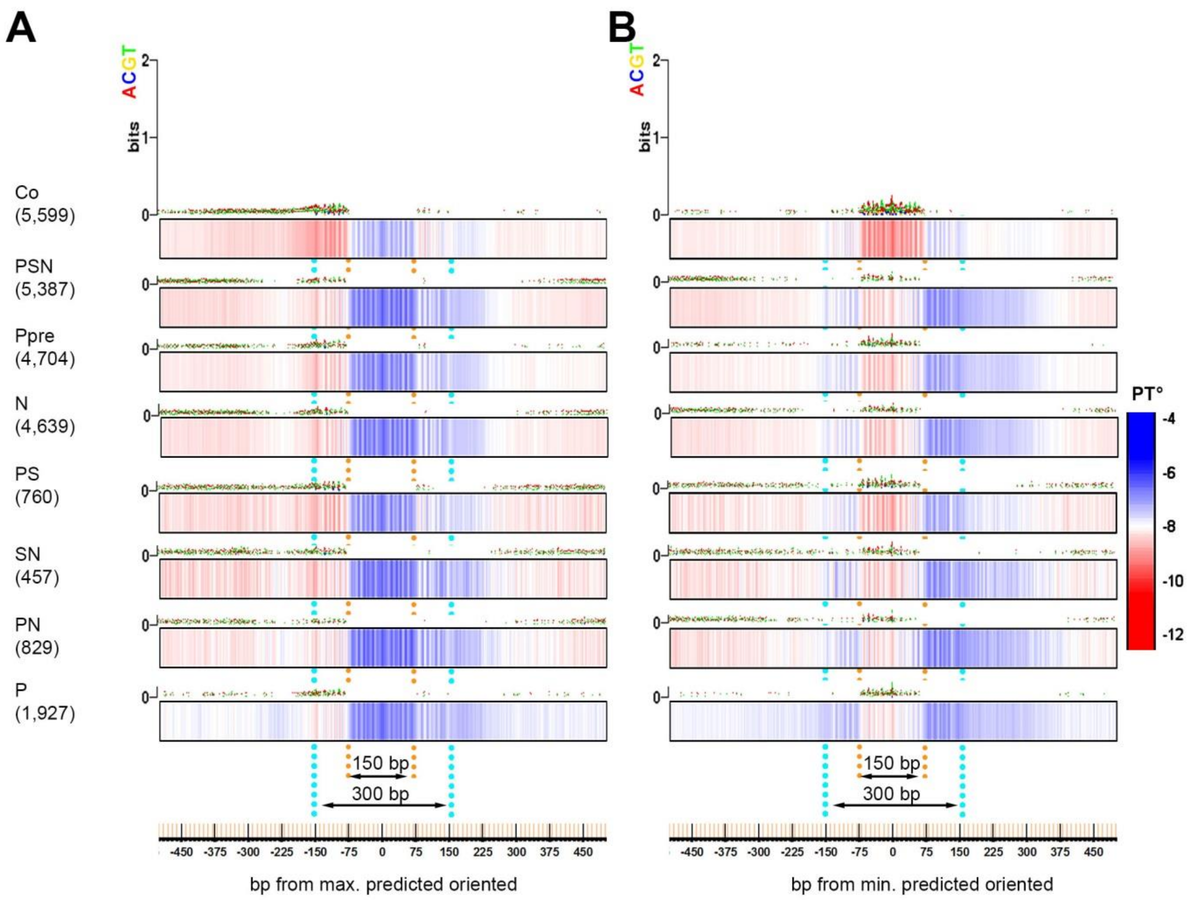

**Fig. S13.**

**PT value sensitively reflects two attributes of DNA affinity to nucleosomes: GC content and dinucleotide rotational periodicity.** Two sequence-based features: increased GC content and AT/GC periodicity, were reported to define high affinity of DNA to nucleosomes. The preference for GC pairs was related to a lower energetic cost required for deformation of the DNA to wrap around the histones (Drew and Travers 1985; Field et al. 2008; Chung and Vingron 2009; Tillo et al. 2010). The nucleosome prediction program of Kaplan et al. (Kaplan et al. 2009) is based to large extent on capturing periodic fluctuations and AT/GC content and does not account for PT or other DNA shape parameters. Here, we compared PT and base composition plots around max. and min. nucleosome occupancy positions (Kaplan et al. 2009). We found that PT value and PT periodic oscillations visualize the sequence properties captured by nucleosome prediction program in a much more sensitive and perhaps direct way than sequence composition itself. Base composition and PT were determined using TFBSshape server (Yang et al. 2014) in 7 TFBS and control groups. The plots were downloaded from the server and contrasted together with the PT scale. (*A*) All genomic regions were centered on max. predicted nucleosome occupancy value [nucmax] (Kaplan et al. 2009) within 320 bp around TFBS. The regions were oriented so that minimum nucleosome prediction value within +/-160 bp around [nucmax] is at the left (5') side. (*B*) All genomic regions were centered on min. predicted nucleosome occupancy value [nucmin] (Kaplan et al. 2009) within 320 bp around TFBS, and oriented so that [nucmin] is at the 5' side from [nucmax] position. Numbers of genomic regions is indicated in parentheses. Note that the Zebrafish genome is AT rich (median GC% = 36.8 (NCBI)), therefore absence of AT/CG bias i.e. around the max. positions indicates the increase in GC content over genomic average.

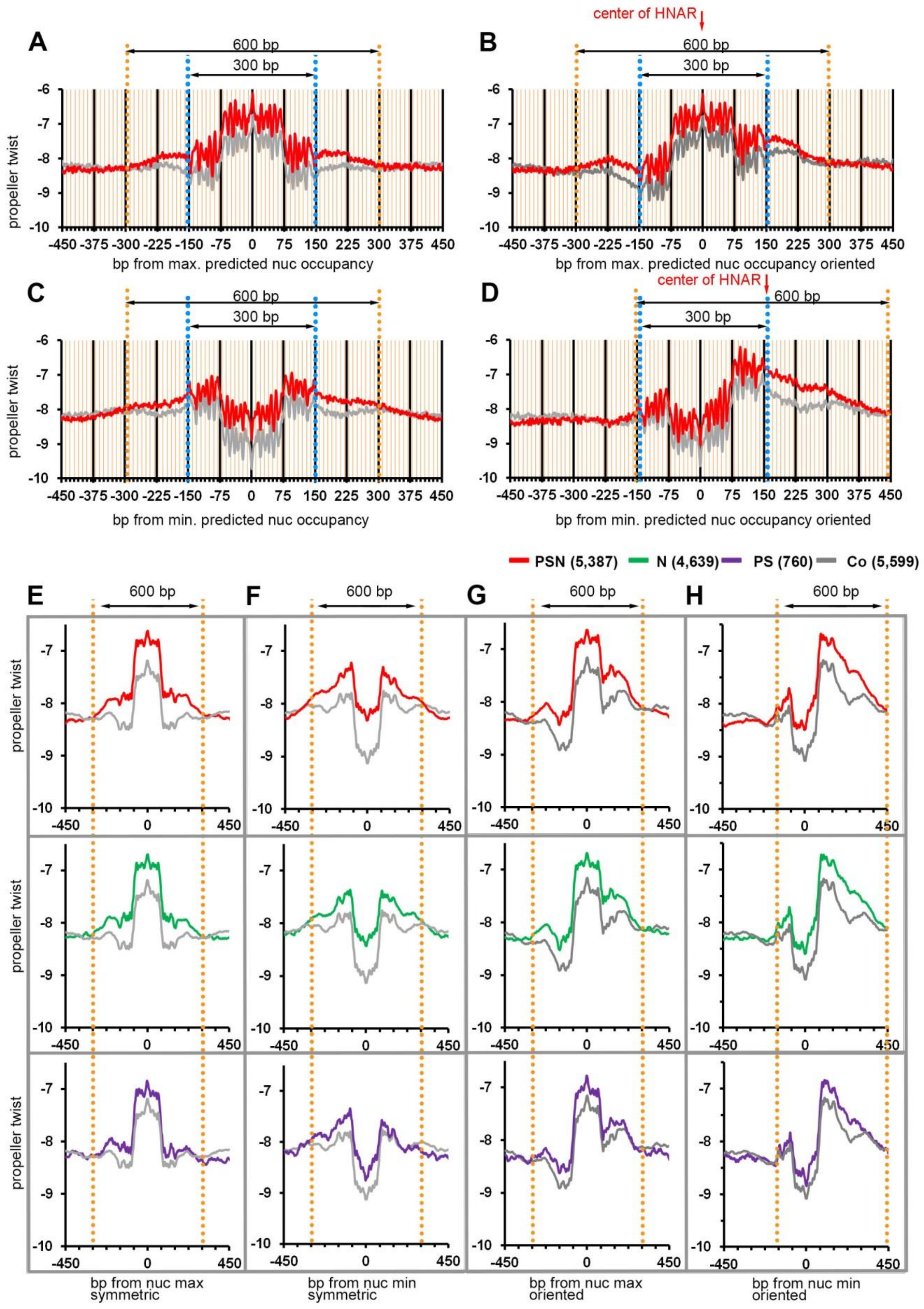

**Fig. S14.**

**Estimation of the length of HNARs PSN, N and PS.** We estimated the length of high PT region by the difference with the control and TFBS in symmetric and oriented plots, as 600 bp (PSN, N) and 450-500 bp (PS). (*A*-*D*) Un-smoothened PT plots and (*E*-*H*) smoothened PT plots (80 bp moving average) of TFBS genomic regions compared to control, centered on predicted dyad [nucmax] or predicted intra-nucleosomal region [nucmin] (Kaplan et al. 2009) within 320 bp around the TFBS, oriented or symmetric as indicated. Average PT value per 1 bp was calculated for – 450 to + 450 bp from central position and plotted in Excel. Blue dotted lines indicate the borders of 300 bp periodic frame. Orange lines indicate the 600 bp when most of TFBS are different from control by higher PT (PSN compared to control). (*B*,*G*) Genomic regions centered on [nucmax] were oriented so that minimum nucleosome prediction value within +/-160 bp around [nucmax] is upstream (at the left side) from [nucmax] position (as in main Fig.3H). (*D*,*H*) Genomic regions centered on [nucmin] were oriented the same way. The center of PSN HNAR is shown by red arrow. Note that [nucmax] is in the center of the HNAR, while [nucmin] is close to the edge of the HNAR in oriented PT plots.

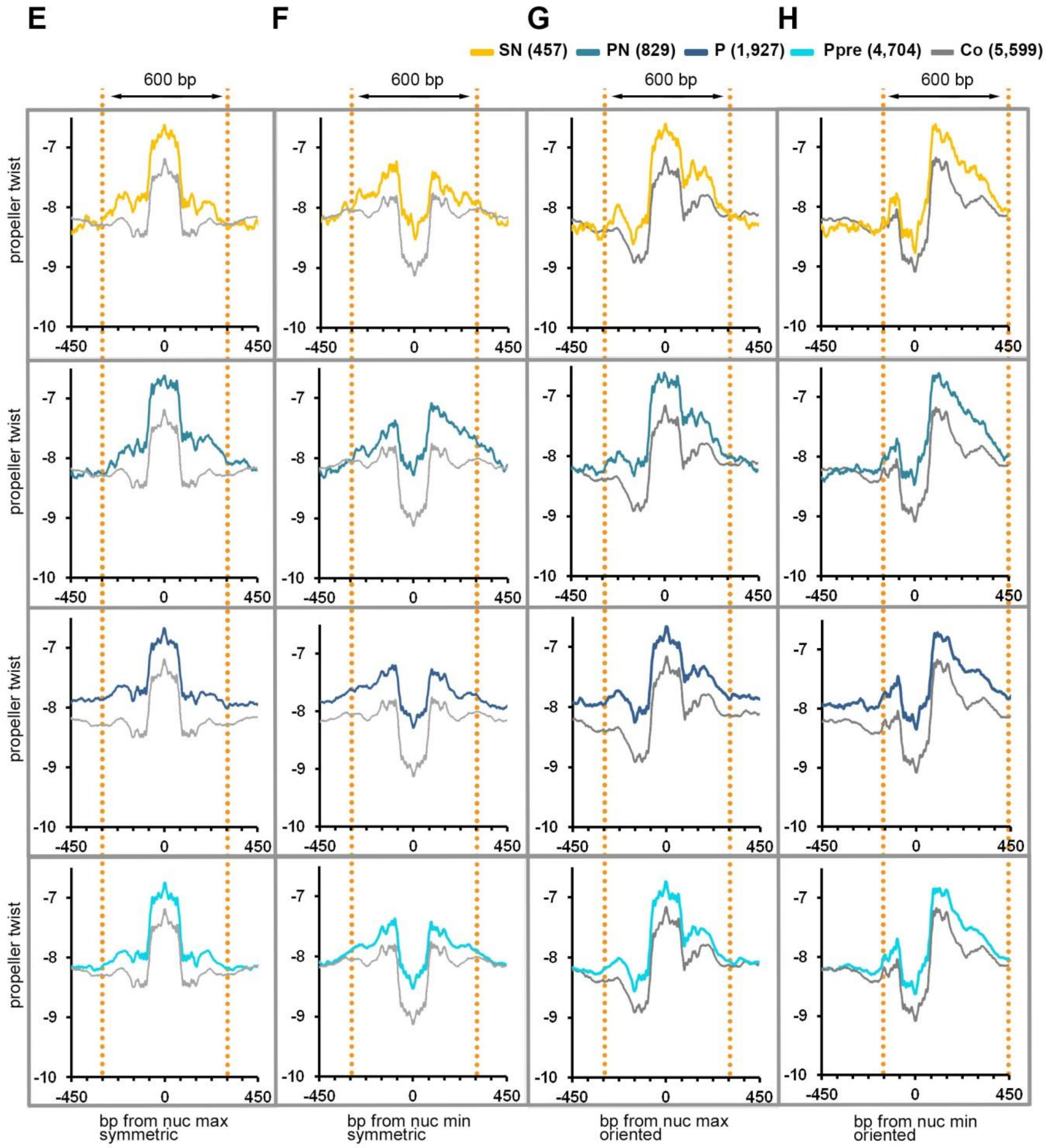

**Fig. S15.**

**Estimation of the length of the HNAR SN, PN, P and Ppre (Fig. S14 continued).** We estimated the length of high PT regions by the difference with the control and TFBS in symmetric and oriented plots as 600-700 bp (Ppre, SN, PN) and >900 bp (P) (*E-H*) see the legend in Fig. S14. Note that in the group of genomic regions, bound by Pou5f3 alone post-ZGA (P), higher PT compared to the control extends beyond +/-450 bp from [nucmin] or [nucmax]. P group was the only TFBS group which did not show enrichment in developmental enhancers, and dinucleotide repeats. We hypothesize that 600 bp bounds of high PT/predicted nucleosome occupancy reflect the potential of DNA sequence with high PT to become an enhancer.

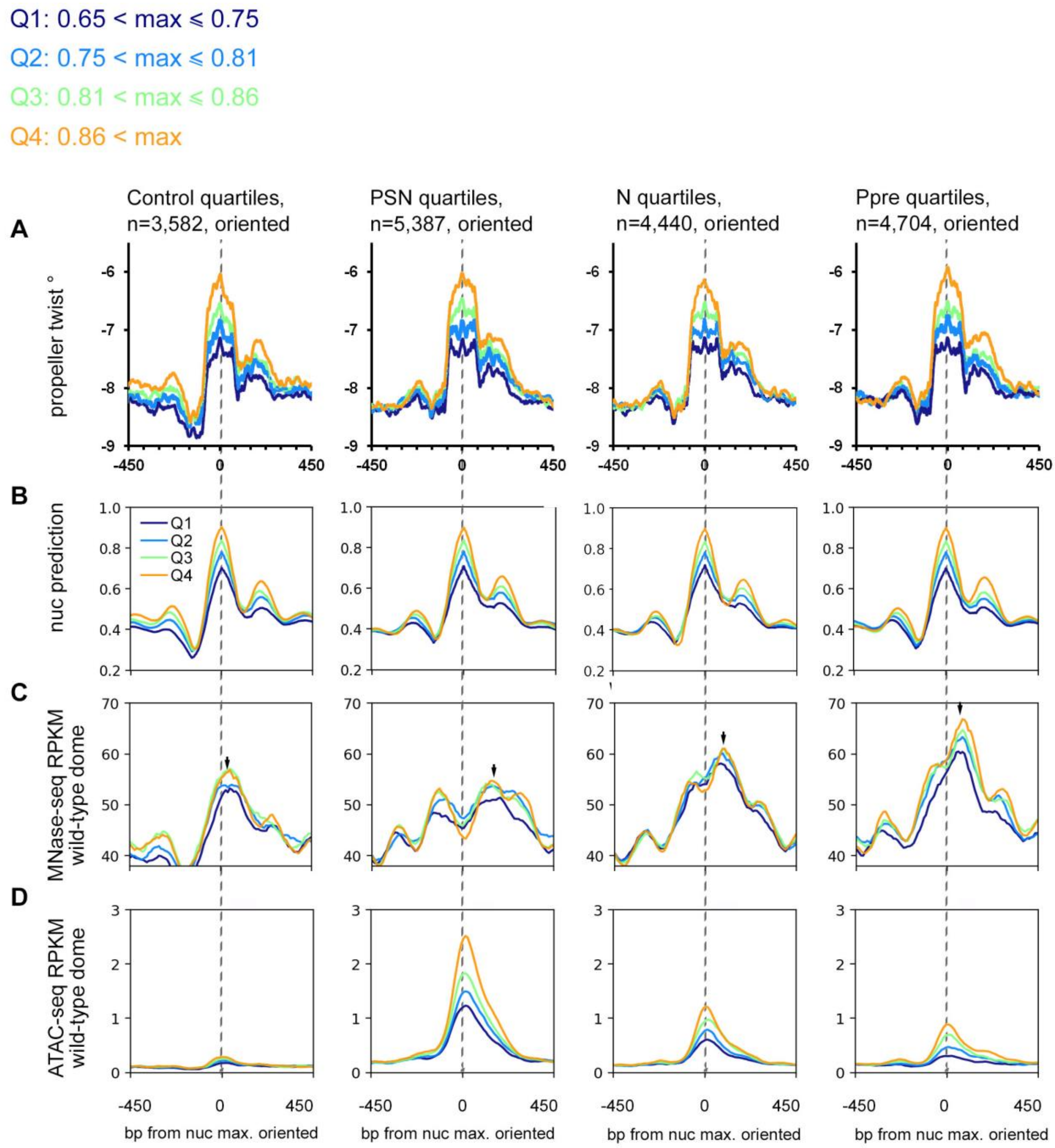

**Fig. S16.**

**Nucleosome destabilization on HNAR centers increases with nucleosome prediction strength in control and all TFBS groups.** 20,747 genomic regions, with max. nucleosome predicted value within 320 bp around the TF binding site/center more than 0.65, were divided to four quartiles according to max. values. Max. values per quartile are shown above. Genomic regions were aligned at [nucmax], and were oriented so that minimum nucleosome prediction value within +/-160 bp is upstream (at the left side) from [nucmax] position. (*A*) Mean PT values per quartile in the indicated groups were calculated in TFBShape server and plotted in Excel. [nucmax] values are listed in Table S1. (*B*) Nucleosome predictions. (*C,D)* ATAC-seq signals, at 256 (*C*) and dome (*D*) stage. Note that ATAC-seq signals peak on maximal nucleosome prediction (gray lines) and increase with predicted nucleosome positioning strength in both pre- and post-ZGA. Gray dotted lines mark HNAR centers.

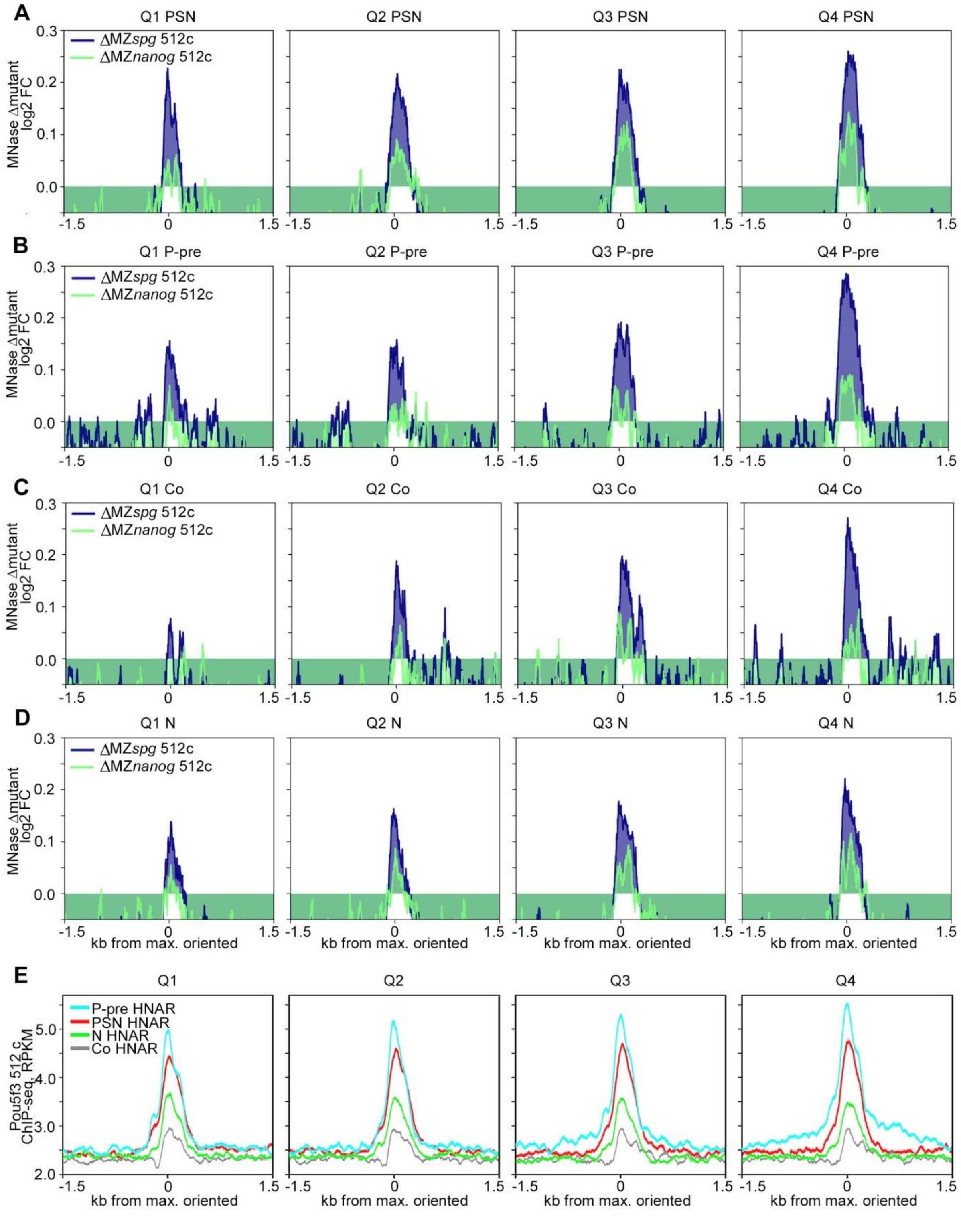

**Fig. S17.**

**Pou5f3 and Nanog non-specifically displace nucleosomes from HNAR centers pre-ZGA (related to Fig. 4 of the main text).** Indicated TFBS regions were ranked by ascending nucleosome prediction score into quartiles Q1-Q4, aligned on [nucmax], and oriented so that minimum nucleosome prediction value within +/-160 bp around [nucmax] is upstream (at the left side) from [nucmax] position (as in main Fig.3H). Plots show the values indicated on top for four quartiles. (*A*-*D*) Nucleosome occupancy changes (Δmut) are localized on HNAR center and increase with nucleosome prediction value. 0 - [nucmax]. (*A*) PSN group (*B*) Ppre group (*C*) Control group (*D*) N group (*E*) ChIP-seq signal of Pou5f3 at 512-cell stage localizes on [nucmax] in all TFBS groups and control.

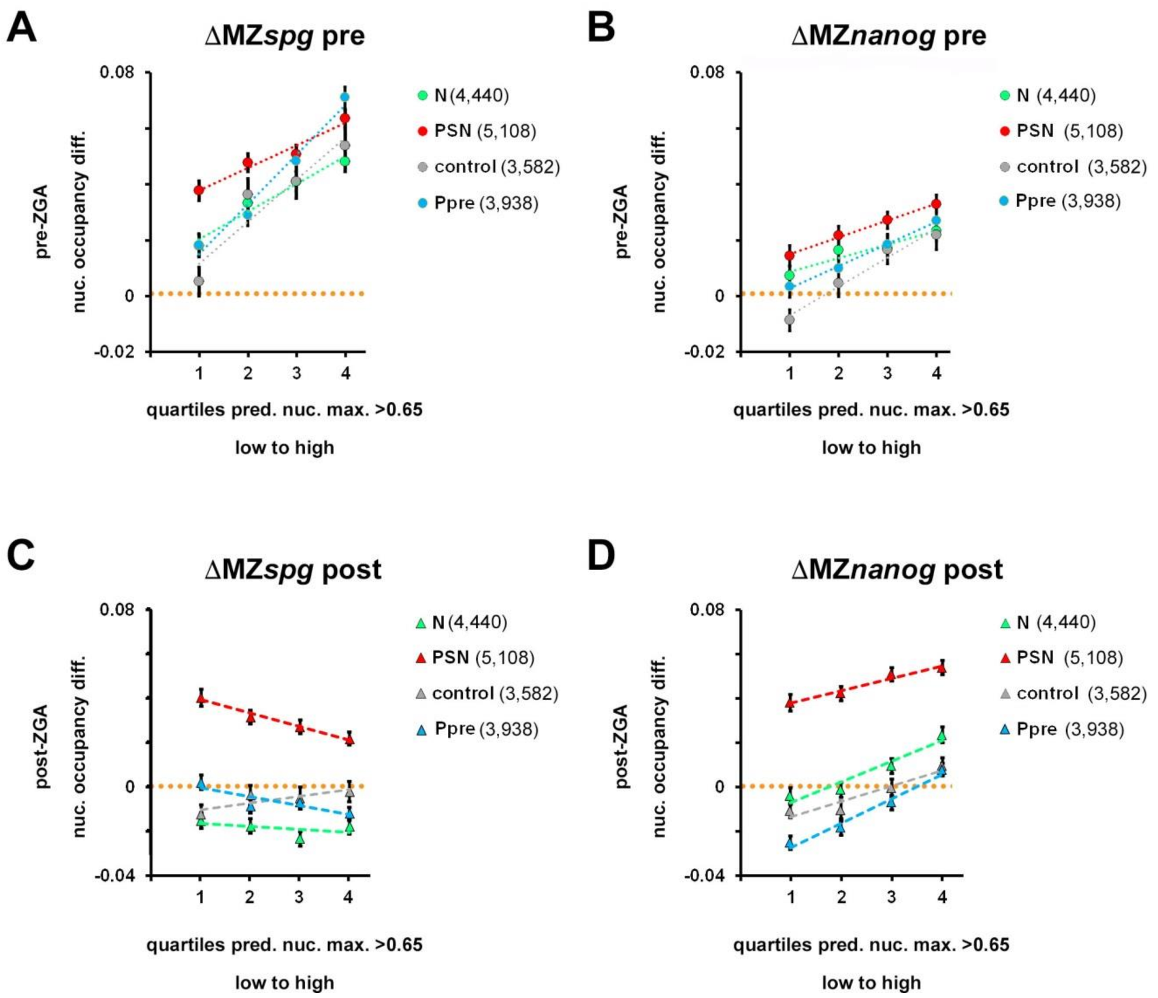

**Fig. S18.**

**ΔMZ*spg* and ΔMZ*nanog* dependence on nucleosome footprint strength in PSN, N, Ppre and control group (related to Fig. 4 of the main text).** Normalized difference between the nucleosome occupancy in the mutant and wild-type (Δmut) for 320 bp region around [nucmax] position was calculated as (rpkm(mut)-rpkm(wt))/(rpkm(mut)+rpkm(wt)). (*A*) pre-ZGA in MZ*spg*, Δmut increases with predicted nucleosome occupancy in all groups. (*B*) Pre-ZGA in MZ*nanog*, Δmut increases with predicted nucleosome occupancy in all groups. (*C*) Post-ZGA in MZ*spg,* Δmut does not depend on predicted nucleosome occupancy in N and control, and decreases with predicted nucleosome occupancy in PSN and Ppre groups. (*D*) Post-ZGA in MZ*nanog,* Δmut increases with predicted nucleosome occupancy in all groups. Note, that however, when compared to pre-ZGA, the Δmut MZ*nanog* threshold is changed: in all groups except PSN, in two lower quartiles Δmut MZ*nanog* is zero or negative.

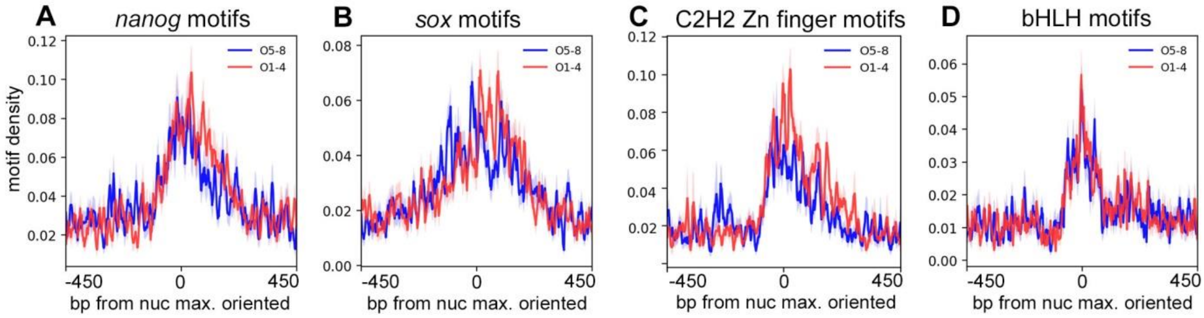

**Fig. S19.**

**Density of centrally located motifs in "open" and "closed" HNARs is similar (related to Fig. 5 of the main text).** PSN group regions ranked into octiles by ΔWT post-pre, aligned on [nucmax] and oriented as in Fig. 3H. Density of the indicated motifs in octiles O1-O4 (red) versus octiles O5-O8 (blue), bp motif per bp sequence.

**Table S3.**
Summary of mapping.

| stage/genotype | total mapped reads |
|---|---|
| Dome_WT | 301153375 |
| Dome_MZ*spg* | 313912467 |
| Dome_MZ*nanog* | 255374362 |
| 512c_WT | 171024005 |
| 512c_MZ*spg* | 156872258 |
| 512c_MZ*nanog* | 144771703 |

**Table S4**.
1-way ANOVA of post to pre ZGA differences nucleosome occupancy at different TF regions in the WT (Fig. S5D), and of mutant to the wild type differences pre- and post-ZGA for MZ*nanog* and MZ*spg* mutants (Fig. S5B,C,E,F).

| Source | DF | Sum of Squares | Mean Square | F Ratio | Prob > F |
|---|---|---|---|---|---|
| | Response: ΔWT(post-pre) | | | Fig.S5 D | |
| TFs_binding | 7 | 33.67 | 4.810 | 147.77 | <.0001 |
| Error | 30506 | 992.91 | 0.033 | | |
| | Response: ΔMut_spg-pre) | | | Fig.S5 B | |
| TFs_binding | 7 | 14.48 | 2.069 | 63.53 | <.0001 |
| Error | 30500 | 993.40 | 0.033 | | |
| | Response: ΔMut_Nanog-pre) | | | Fig.S5 C | |
| TFs_binding | 7 | 5.43 | 0.776 | 28.48 | <.0001 |
| Error | 30495 | 830.96 | 0.027 | | |
| | Response: ΔMut_spg-post) | | | Fig.S5 E | |
| TFs_binding | 7 | 14.69 | 2.099 | 98.14 | <.0001 |
| Error | 30507 | 652.44 | 0.021 | | |
| | Response: ΔMut_Nanog-post) | | | Fig.S5 F | |
| TFs_binding | 7 | 34.35 | 4.907 | 219.33 | <.0001 |
| Error | 30508 | 682.49 | 0.022 | | |

**Table S5.**
1-way ANOVA of mutant-WT nucleosome occupancy differences pre-ZGA and post-ZGA (main Fig. 2 E) at TF-binding regions with different number of non-overlapping TF binding sites. Control values are not included into this analysis. Colors correspond to those on Fig. 2E.

| Source | DF | Sum of Squares | Mean Square | F Ratio | Prob > F |
|---|---|---|---|---|---|
| | Response: [delta]Mut_Nanog-pre) | | | | |
| N(nanog motifs) at N regions | 4 | 0.10 | 0.024 | 1.08 | 0.37 |
| Error | 5448 | 120.46 | 0.022 | | |
| N(nanog motifs) at PSN regions | 4 | 0.08 | 0.021 | 0.94 | 0.44 |
| Error | 6327 | 140.12 | 0.022 | | |
| | Response: [delta]Mut_Nanog-post) | | | | |
| N(nanog motifs) at N regions | 4 | 0.02 | 0.004 | 0.31 | 0.87 |
| Error | 5448 | 76.96 | 0.014 | | |
| N(nanog motifs) at PSN regions | 4 | 1.86 | 0.466 | 21.65 | 9.10E-18 |
| Error | 6327 | 136.04 | 0.022 | | |
| | Response: [delta]Mut_spg-pre) | | | | |
| N(pou:sox motifs) at PS regions | 4 | 0.04 | 0.010 | 0.32 | 0.87 |
| Error | 941 | 29.73 | 0.032 | | |
| N(pou:sox motifs) at PSN regions | 4 | 0.36 | 0.089 | 4.05 | 0.0028 |
| Error | 6327 | 139.61 | 0.022 | | |
| | Response: [delta]Mut_spg-post) | | | | |
| N(pou:sox motifs) at PS regions | 4 | 0.25 | 0.063 | 3.58 | 0.0067 |
| Error | 941 | 16.66 | 0.018 | | |
| N(pou:sox motifs) at PSN regions | 4 | 7.81 | 1.953 | 103.46 | 1.74E-85 |
| Error | 6327 | 119.44 | 0.019 | | |

**Table S6.**
T-tests of the difference between nucleosome occupancy (Δmut) for the TF bound regions lacking motifs and that in randomly chosen control regions, pre-ZGA and post-ZGA (main Fig. 2E). Colors correspond to those on the figure.

| | Mutant=MZ*nanog* | | | | Mutant=MZ*spg* | | | |
|---|---|---|---|---|---|---|---|---|
| | Δmut_pre | | Δmut_post | | Δmut_pre | | Δmut_post | |
| | t | p | t | p | t | p | t | p |
| Control vs. PSN *pou:sox* motifs=0 | 8.82 | 6.68E-19 | 16.09 | 1.71E-57 | 14.11 | 9.59E-45 | 2.05 | 0.04 |
| Control vs. PS *pou:sox* motifs=0 | | | | | 2.92 | 0.0035 | -0.31 | 0.76 |
| Control vs. N *nanog* motifs=0 | 5.93 | 3.10E-09 | 4.94 | 7.81E-07 | | | | |

**Table S7 and S8.**
1-way ANOVA of the differences between [nucmax] and [nucmin] among TF binding groups. Related to Fig. S10.

| | | | | | |
|---|---|---|---|---|---|
| | Response: | nucmax | | | |
| Source | DF | Sum of Squares | Mean Square | F Ratio | Prob > F |
| TFs_binding_group | 7 | 61.49 | 8.78 | 521.15 | 0 |
| Error | 24294 | 409.47 | 0.02 | | |
| | Response: | nucmin | | | |
| Source | | | | | |
| TFs_binding_group | 7 | 60.50 | 8.64 | 523.81 | 0 |
| Error | 24294 | 400.82 | 0.02 | | |

**Table S9**.
Linear regression of ATAC-seq signals on the predicted nucleosome occupancy (truncated by removal of estimates <0.65) at different developmental stages.

| Developmental stage | $R^2$ | Intercept | Slope | P |
|---|---|---|---|---|
| 64 cells | 0.034 | -0.150 | 0.462 | <.0001 |
| 256 cells | 0.068 | -0.220 | 0.562 | <.0001 |
| 1K | 0.046 | -0.375 | 0.887 | <.0001 |
| oblong | 0.064 | -0.788 | 1.505 | <.0001 |
| dome | 0.045 | -1.160 | 2.155 | <.0001 |

**Table S10**.

1-way ANOVA of mutant-WT nucleosome occupancy differences pre- and post-ZGA at regions that fall into 1st through 4th quartiles of predicted nucleosome occupancy (main Fig. 4G). Colors correspond to those on the figure.

| Source | DF | Sum of Squares | Mean Square | F Ratio | Prob > F |
|---|---|---|---|---|---|
| Response: Δmut MZ*nanog* pre-ZGA | | | | | |
| [nucmax] quartiles | 3 | 1.238 | 0.413 | 16.69 | 7.92E-11 |
| Error | 20739 | 512.538 | 0.025 | | |
| Response: Δmut MZ*spg* pre-ZGA | | | | | |
| [nucmax] quartiles | 3 | 3.547 | 1.182 | 42.61 | 1.93E-27 |
| Error | 20743 | 575.578 | 0.028 | | |
| Response: Δmut MZ*nanog* post-ZGA | | | | | |
| [nucmax] quartiles | 3 | 2.403 | 0.801 | 38.60 | 7.28E-25 |
| Error | 20743 | 430.485 | 0.021 | | |
| Response: Δmut MZ*spg* post-ZGA | | | | | |
| [nucmax] quartiles | 3 | 0.156 | 0.052 | 2.77 | 0.040 |
| Error | 20743 | 389.867 | 0.019 | | |

**Table S1. (separate file)**

Genomic regions used in this study, nucmax and nucmin positions and values, strand for oriented plots.

**Table S2. (separate file)**

Positional Weight Matrices for the motifs found in this study.
